# The Eukaryotic homology search complex distorts donor DNA structure to probe for homology

**DOI:** 10.1101/2025.08.28.672940

**Authors:** Mitchell V. Woodhouse, Jingyi Hu, Meiling Wu, Jin Qian, James T. Inman, Michelle D. Wang, J. Brooks Crickard

## Abstract

Homologous recombination (HR) is a DNA double-strand break repair pathway that facilitates genetic exchange and protects damaged replication forks during DNA synthesis. As a template-based repair process, the successful repair of a double-strand break depends on locating suitable homology from a donor DNA sequence elsewhere in the genome. In eukaryotes, Rad51 catalyzes the homology search in coordination with the ATP-dependent motor protein Rad54. The mechanism by which these two proteins regulate forces on dsDNA substrates during homology search remains unknown. Here, we have utilized single-molecule magnetic tweezers and optical trapping methods to monitor remodeling of the DNA template during the homology search. We find that the activity of Rad51 and Rad54 remodels the donor DNA substrate to control the association and dissociation of Rad51-ssDNA filaments in the absence of DNA homology. This mechanism occurs through the application of both linear (tension) and rotational (torsion) forces on the donor DNA. Finally, failure of Rad54 to act processively disrupts target selection *in vivo*. This study provides a basic understanding of how motorized homology search manipulates the donor DNA during the search for a suitable repair template.

**Significance Statement:** Homologous recombination (HR) is a double-strand DNA break repair pathway that utilizes a template-based target search process to locate a suitable homologous DNA sequence in the genome, thereby initiating DNA repair. Called the homology search, in eukaryotes, this process is carried out by the RecA family member Rad51. During the homology search, Rad51 collaborates with the motor protein Rad54 to identify and interrogate homologous DNA sequences within the genome. In this study, we have measured the forces applied by the combination of Rad51 and Rad54 to the donor DNA duplex. These measurements reveal a coordinated effort by these motor proteins to remodel donor DNA to probe for homology, shedding new light on how template-based homology searches interrogate the DNA strands.

## Main Text

Homologous recombination (HR) is a universally conserved template-based DNA double-strand break repair (DSBR) pathway that requires a recipient ssDNA to locate a matching donor DNA elsewhere in the genome (1–3). Finding a suitable donor occurs through a homology search process. This systematic effort initially searches sequences local to the break site and then can extend to more distal regions until homology is located (4–6). Members of the RecA recombinase family control the selection and stabilization of the donor DNA sequence (7–16).

The RecA family of recombinases is conserved in all domains of life. The basic biochemical mechanism of these proteins involves the formation of a filament on the single-stranded recipient DNA, where it binds and stabilizes the ssDNA at 1.5 times the contour length of B-form DNA (10,17). The three-nucleotide spacing of individual protomers in this filament enables the coordination of two DNA-binding sites that can bind and stretch the donor duplex, promoting the kinetic sampling of base pairs (12). Binding site I is composed of DNA-binding loops from distinct RecA protomers that allow the unstacking of bases from the parent donor duplex (10,17,18) and promote base flipping to enhance contacts between the incoming recipient and the donor DNA. A second DNA-binding site, DNA-binding site II, interacts with the non-homologous strand of DNA, stabilizing the separation of the two parent strands (19). The two binding sites coordinate during the homology search to actively probe donor DNA sequences. A minimum of 8 paired nucleotides is required for stable kinetic sampling (8,12,14). A spacing of four recombinase protomers, resulting in 12 paired nucleotides, is optimal for stable sequence selection (18,20). Further recipient donor pairing is considered a strand exchange reaction, leading to a displacement loop (D-loop).

Both DNA binding sites I and II favor binding to ssDNA (20), making dsDNA a poor substrate for Rad51-ssDNA filaments. Partial separation of DNA strands can promote the binding of recombinase filaments in the absence of DNA sequence homology, as the duplex DNA begins to resemble single-stranded DNA (ssDNA). The underwinding of DNA can lead to partial strand separation by reducing the number of bases per turn of the regular B-form helix. Mechanically, this can be achieved by linear stretching, which involves adding tension, or by rotating the DNA around the superhelical axis, which adds torsion (21–23). It has been experimentally determined that the addition of forces by stretching (tension) (24,25) or rotating (torsion) (20,26) can improve the binding of recombinase-ssDNA filaments to the donor DNA in the absence of DNA sequence homology. The superhelical density of DNA *in vivo* is generally underwound, improving recombinase filament binding. However, this is not uniform, which makes regulation of DNA topology a crucial feature in controlling the binding of recombinase filaments to the donor DNA during the homology search. The factors that aid recombinases in homology search may impact early DNA sequence recognition by providing assisting forces to regulate topology, but the mechanism behind this is unclear.

In eukaryotes, the ATP-dependent translocase Rad54 aids Rad51 during the homology search and strand exchange (27–32). Rad54 is related to chromatin remodeling enzymes of the Snf2 family (33) and has also demonstrated nucleosome remodeling activity (29,34–36). The fundamental activity of Rad54 is to physically move along dsDNA, tracking the minor groove in the 3’ to 5’ direction (37–39). Rad54 movement can remodel the DNA by stretching, bending, or twisting the DNA helix. The magnitude of these remodeling events has not been measured. However, they can result in the removal of other proteins from DNA or the addition of negative twist to the DNA’s superhelical axis. The resulting accumulation of supercoils in the DNA requires the DNA to be topologically locked. Recent evidence has emerged that negative supercoils may accumulate even on linear DNA. The mechanism behind this is unknown.

Rad51 enhances the ATP hydrolysis and translocation activity of Rad54, and together, they can form a homology search complex (28,29,40–42). The interaction between Rad54 and Rad51 occurs through Rad54’s intrinsically disordered N-terminal domain (43,44). This region can interact with Rad51 and form a multimerization domain with other Rad54 molecules. Removal or replacement of the N-terminal domain renders *Saccharomyces cerevisiae* cells sensitive to DNA damaging agents, comparable to the *rad54Δ* strains (43–46). Functional activities of Rad54 include the disruption of Rad51 filaments bound to dsDNA (47–50) and the remodeling of dsDNA during the homology search (29,51,52). The addition of Rad54 to Rad51 filaments introduces a 1D translocation-based search, which can accelerate the identification of homologous DNA (29).

The application of forces can promote the binding of Rad51 filaments to donor dsDNA. Rad54 is a motor capable of adding these forces and forms a complex with Rad51 during the homology search. In this study, we investigated how forces added by Rad54 influence the homology search activity of Rad51-ssDNA filaments before the identification of homologous regions. We find that Rad54 can form isolated looped regions on the donor DNA, promoting loop extrusion. Isolation allows the addition of torsional stress to the donor DNA, resulting in the storage of negative turns in the DNA. Rad54 also added linear tension to the DNA. We hypothesize that the combination of forces provided by Rad54 stabilizes the binding of Rad51-ssDNA to the donor duplex by underwinding the DNA, thereby catalyzing the base sampling and the homology search process. Importantly, stabilization is reversible through the hydrolysis of ATP. We also demonstrate that mutant forms of Rad54 defective in the homology search fail to form displacement loops *in vivo*. Together, our data provide a novel model for how remodeling of donor DNA structure can catalyze the homology search.

## Results

### PSC activity is dependent on the tension placed on the donor DNA

By tracking the minor groove of dsDNA, Rad54 introduces helical twist to the DNA backbone (39,53). The torsional stress generated by translocation accumulates only if it occurs in a topologically locked region of donor DNA (29,51,54–56). A locked region can form when a group of proteins forms multiple points of contact with the DNA. In this scenario, one contact can pump DNA into an isolated loop, while another serves as an anchor. A consequence of this mechanism would be the apparent ability to extrude loops, resulting in DNA compaction. Previous reports have shown that the Rad54 paralog Rdh54 can move along dsDNA by loop extrusion and directional translocation (57). Rad54 or the combination of Rad54, Rad51, and ssDNA, known as the presynaptic complex (PSC) (29,30), has only been observed to promote translocation, not loop extrusion.

Extruded loops are regulated by the tension applied to the DNA. This is due to the minimal distances required to bend the DNA and form an initial loop. At lower tension, dsDNA is more flexible and accessible to multiple protein-DNA connections, which can result in an isolated region of DNA (58). We hypothesized that the PSC would be more likely to compact DNA or extrude loops at lower tension if this is a mechanism by which it acts. To measure this, we used a dual optical trap with confocal microscopy to monitor the binding of the PSC to dsDNA. We formed the PSC with GFP-Rad54 in combination with Atto647N 90-mer ssDNA bound by Rad51, a strategy we have employed previously to monitor active PSCs (29,30,45). Yeast Rad51-ssDNA does not appreciably bind to dsDNA, unless it interacts with Rad54, the DNA is significantly underwound, or the concentration is sufficiently high (59). Under the conditions used here, Rad54 is required for Rad51-ssDNA binding to dsDNA.

Application of a constant force via a force clamp controls the amount of tension on the donor dsDNA, allowing us to test the hypothesis that the PSC could compact DNA and/or promote loop extrusion through multiple points of contact. We measured the activity of the PSC at forces of 0.5, 1.0, 2.0, and 5 pN (**Figure 1A**). The PSC primarily compacted DNA at 0.5 and 1.0 pN (**Figure 1BC**) and moved along the DNA without compacting at 2.0 and 5.0 pN (**Figure 1C**). These activities were dependent on the hydrolysis of ATP. The rates of compaction and movement were comparable, with the rate of compaction at 0.5 pN being 253 ± 193 bp/sec. The measured movement rate at 2 pN was 272.6 ± 133 bp/sec and 357 ± 195 at 5 pN (**Figure 1D**), suggesting that the motor was not inactivated at higher forces. These data support the idea that the PSC can perform loop extrusion of the donor DNA, resulting in compaction.

**Figure 1:**
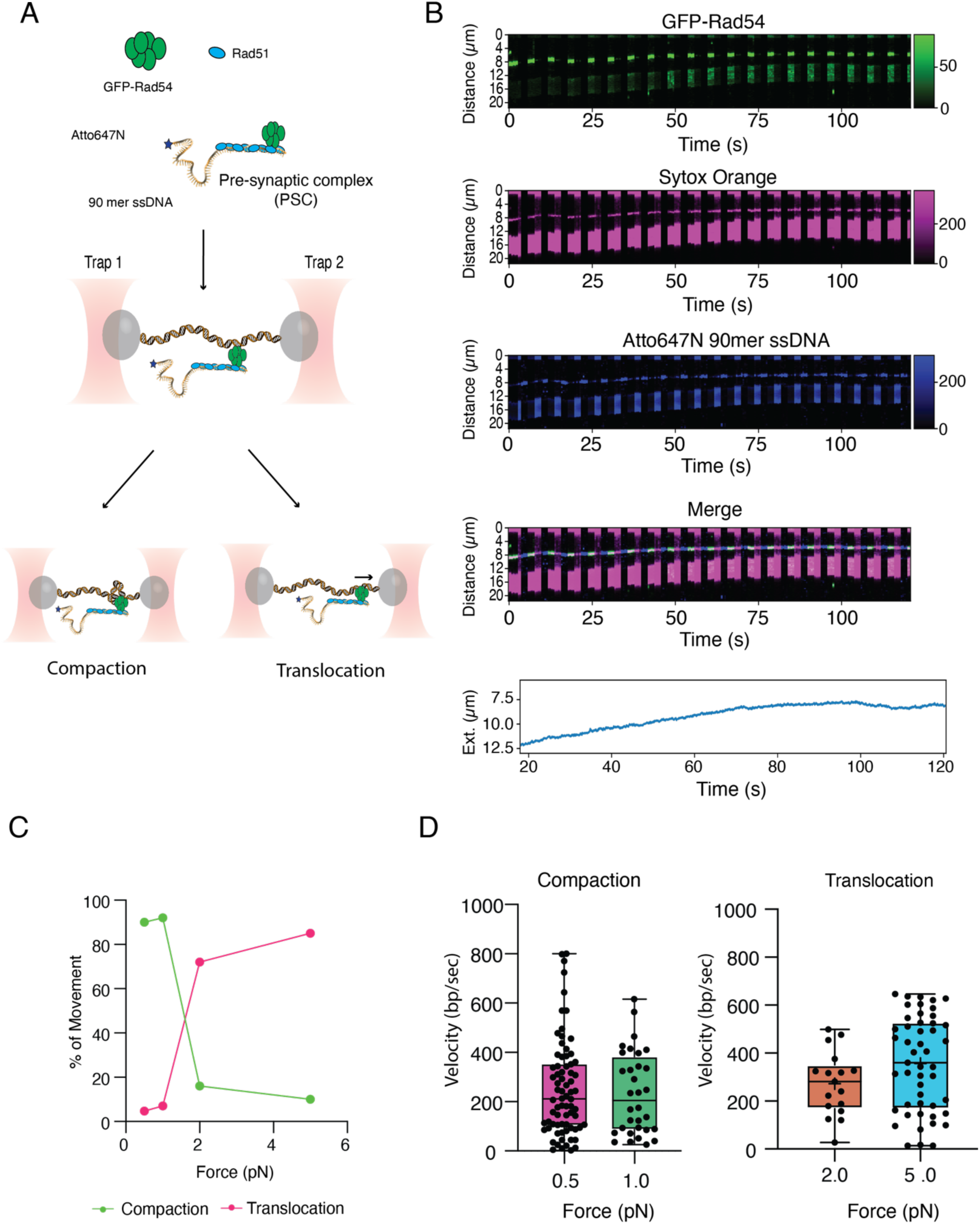
The PSC switches from compaction to translocation at higher force. **(A).** Cartoon diagram illustrating the design to measure the real-time activity of the PSC **(B).** Representative kymographs for PSC activity at 0.5 pN. GFP-Rad54 (Top), Sytox Orange donor DNA (Top middle), Atto647N 90-mer-ssDNA (Bottom middle), and merged (Bottom). Below the kymographs is an extension curve taken during the experiment. **(C)**. Graph representing the percentage of molecules that undergo compaction at 0.5 (N=19/20), 1.0 (N=13/14), 2.0 (N=3/16) and 5.0 (N=2/17) or translocation at 0.5 (N=1/20), 1.0 (N=1/14), 2.0 (N=13/16), and 5.0 (N=17/19). **(D).** Box plots illustrate the compaction rate at 0.5 (N=74) and 1.0 (N=40) pN (left) and the translocation rate at 2.0 (N=20) and 5.0 (N=49) pN (right). The solid line illustrates the mean, and the error bars represent the range of the data.

We tested whether loop isolation occurred through a single contact or multiple points of contact. To measure this, the DNA was allowed to compact and then re-extended by separating the optical trap at a constant rate (**Figure 2AB**). If compaction was due to multiple points of contact, then re-extension of the compacted DNA should result in a sawtooth pattern as contact points are disrupted (60,61). Disruption of contacts could occur through direct loss of the physical interaction between the protein and DNA, or the sliding of the protein along the DNA, resulting in a slip-stick mechanism. For simplicity, we will view both mechanisms as potential disruption events. When we pulled on the compacted DNA molecules at 0.5 and 1.0 pN, they displayed sawtooth patterns (**Figure 2C**). As expected, this behavior was not observed at higher forces because there was no compaction, and all molecules followed a theoretical force extension (FE) curve consistent with B-form DNA (**Figure 2D**).

**Figure 2:**
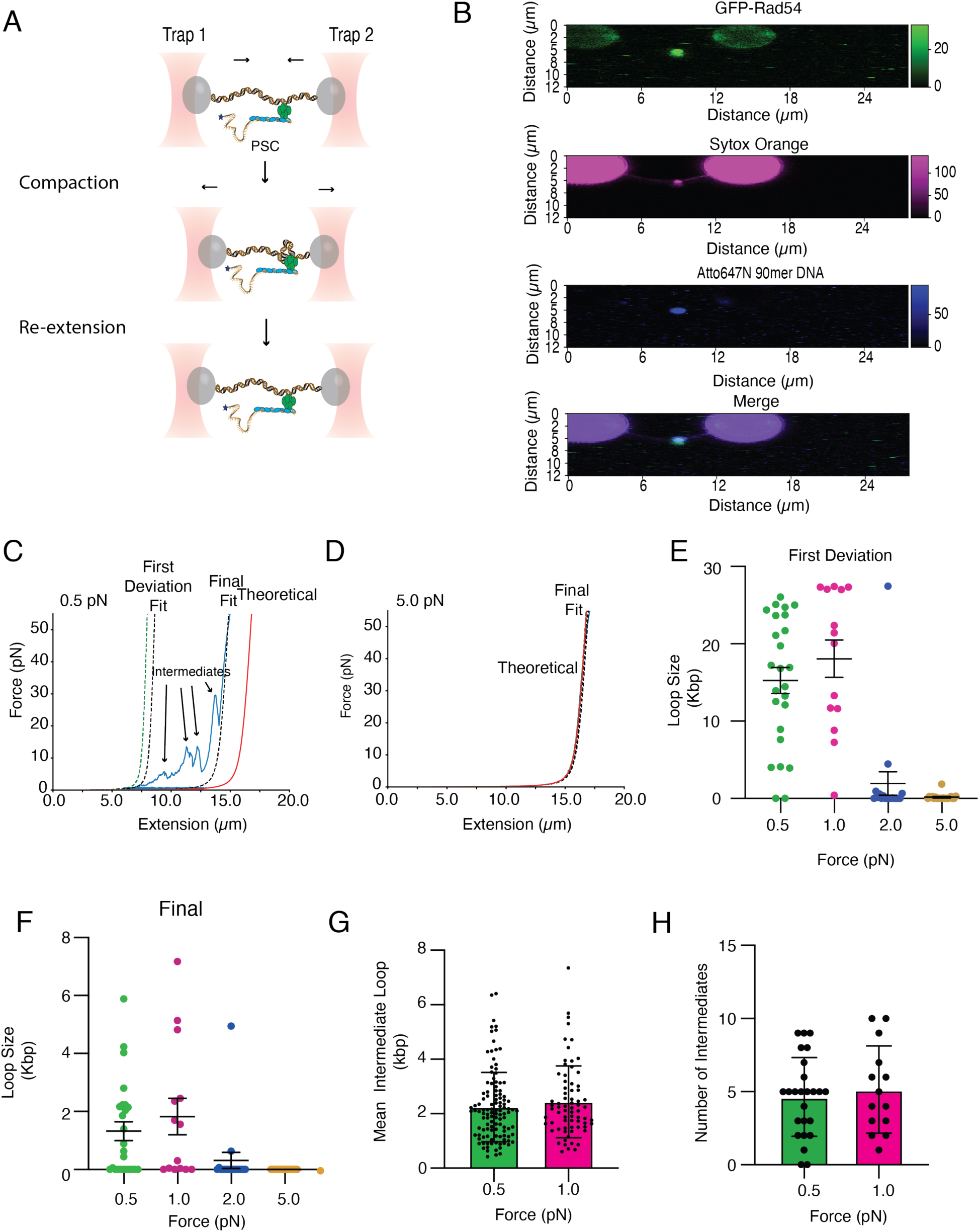
The PSC compacts donor DNA in a force-dependent manner. **(A).** Cartoon schematic diagram of the optical trapping experiment designed to determine if compaction generates multiple points of contact. **(B).** Representative widefield image of GFP-Rad54 (Top), Sytox Orange donor DNA (Top middle), Atto647N 90-mer ssDNA (Bottom middle), and merged (Bottom). **(C).** Force extension curve for the re-extension of a DNA molecule compacted by the PSC at 0.5 pN. The theoretical line is in red, and the data is in blue. The dashed lines represent fits to the traces. The green dashed line is the initial point of deviation of the theoretical FE curve **(D).** Force extension curve for the re-extension of a DNA molecule compacted by the PSC at 5.0 pN. The theoretical line is in red, and the data is in blue. The dashed lines represent fits to the traces. **(E).** Graph illustrating the degree of DNA compaction at first deviation from theoretical for 0.5 (N=25), 1.0 (N=14), 2.0 (N=18), and 5.0 (N=20) pN. The dots represent the mean, and the error bars represent the 95% confidence interval of the data. **(F).** Graph illustrating the degree of DNA compaction at final deviation from theoretical for 0.5 (N=25), 1.0 (N=14), 2.0 (N=18), and 5.0 (N=18) pN. The dots represent the mean, and the error bars represent the 95% confidence interval of the data. **(G).** Graph representing the mean intermediate loop size at 0.5 (N=115) and 1.0 (N=71) pN. The bars represent the mean, and the error bars represent the standard deviation of the experiment. **(H).** Graph representing the number of intermediates per tether at 0.5 (N=25) and 1.0 (N=14) pN. The bar represents the mean, and the error bars represent the standard deviation of the experiment.

The initial amount of compacted DNA is defined as the first deviation from the theoretical FE curve and represents the initial amount of compacted DNA (**Figure 2C**). We continued pulling on the DNA past this point and observed multiple intermediate deviations from the theoretical extension curve. These intermediates reflect numerous contact points between the PSC and the DNA (**Figure 2C**). Structures formed at 0.5 and 1.0 pN of force initially constrained around 15 kbp of DNA. The final size of constrained DNA was around 2 kbp (**Figure 2EF**). The mean of the DNA isolated in intermediates was approximately 2-2.5 kbp per intermediate at both 0.5 and 1.0 pN (**Figure 2G**). We observed a mean contact number of 4-5 intermediates per molecule (**Figure 2H**). It should be noted that due to the small size of the isolated loops, we were unable to visualize them directly. However, based on these data, we conclude that the PSC can isolate individual donor DNA loops of approximately 2 kilobases in length.

### Rad54 applies a significant force during translocation

Previous studies have investigated the effect of tension on Rad51-ssDNA binding to a dsDNA substrate. These studies found that adding tension in the 3’ to 5’ direction of the donor dsDNA could stabilize binding of Rad51-ssDNA or RecA-ssDNA in the absence of DNA sequence homology. When tension was released, the Rad51-ssDNA filaments dissociated (24,25). We reasoned that Rad54 may exert a comparable force on the donor DNA, which could help stabilize Rad51-ssDNA binding to the donor DNA in the absence of homology. We tested the force produced by the PSC during translocation and binding to the donor DNA. Rad54 forms a multimer. Multimeric DNA motors can act in series or parallel depending on the structural organization of the motor. Motors existing in these conformations can make measuring an isolated motor or unit complicated.

Therefore, to isolate the force applied by an individual motor or unit, the PSC was loaded at extensions of 6, 8, 10, 12, and 14 µm of the donor DNA. These values represent 36, 49, 61, 61, 73, and 81% of the contour length of lambda DNA (16.4 µm). By loading the PSC at different extensions, a trend should be observable as the number of contacts between the PSC and donor DNA is reduced. To measure the force during translocation, the beads were moved to a 12.5 µm extension after loading (**Figure 3A**). The extension was maintained to prevent compaction of the beads, allowing measurement of the force output as the PSC moves along the donor DNA (**Figure 3BC**). The force output was not constant during translocation, and we measured the maximum force outputs during individual translocation events (**Figure 3BC**). PSCs loaded at 6 and 8 µm had mean max force outputs of 40 and 23 pN (**Figure 3D**). This force was significantly greater than the mean max forces measured after loading the complex at 10, 12, and 14 µm, which were ∼6-8 pN (**Figure 3D**). These values were greater than those observed for Rad54 alone at 6 and 14 µm loads (**Figure 3D**) and were dependent on ATP hydrolysis (**Figure 3D**). From this, we conclude that the PSC can produce a significant force when translocating on the donor dsDNA.

**Figure 3:**
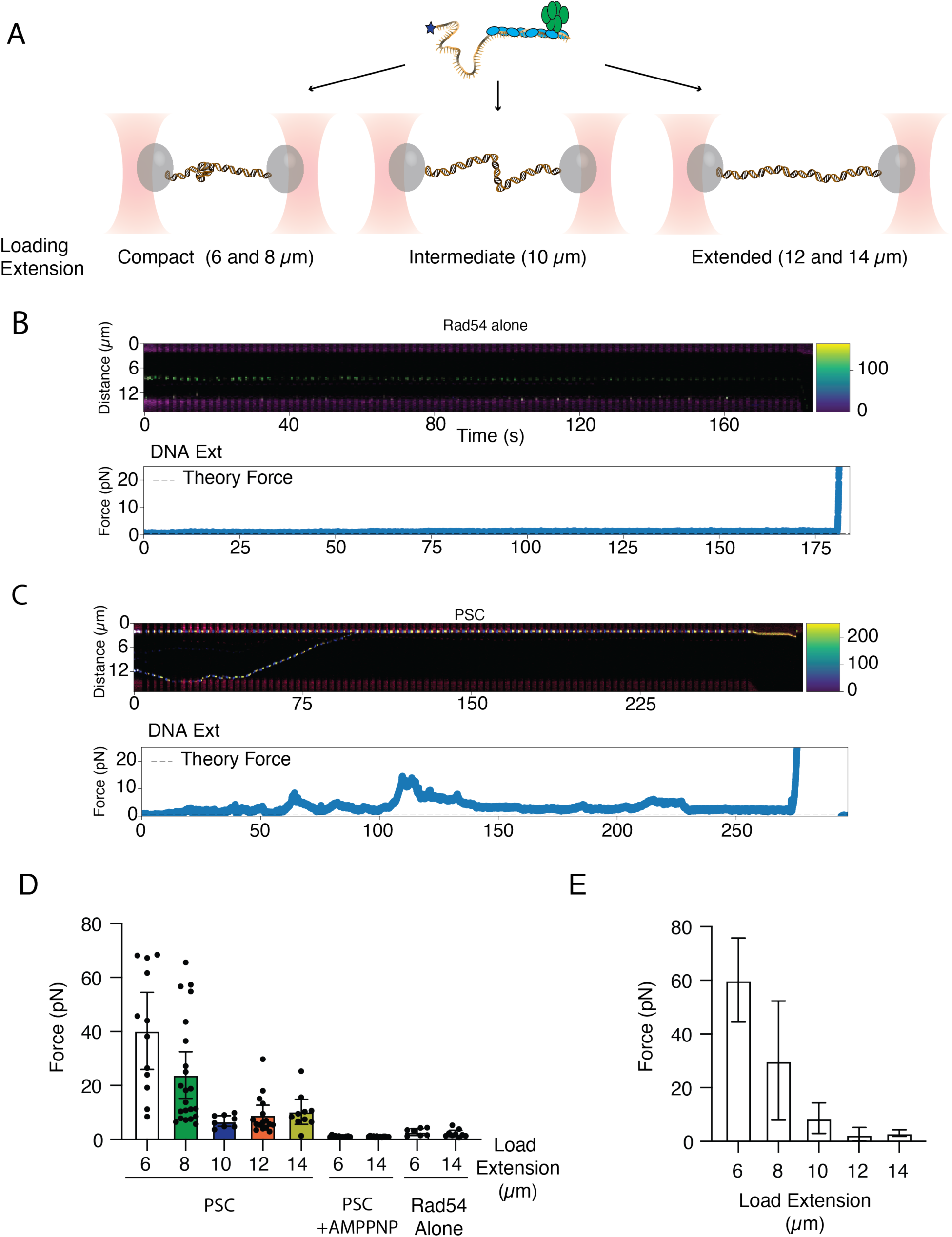
PSC generates a significant force during translocation. **(A).** Cartoon illustration of extension loading experiment. The PSC was loaded at different DNA extensions. **(B).** Representative kymograph and Force trace for Rad54 alone. (C). Representative kymograph and Force trace for the PSC. **(D).** Maximum Force output during activity measurements for the PSC +ATP at 6 (N=12), 8 (N=22),10 (N=8),12 (N=16),14 (N=10) loading extensions, PSC +AMPPMP at 6 (N=12) and 14 (N=13) µm loading extensions, and Rad54 at 6 (N=7) and 14 (N=9) loading extensions. The bars represent the mean, and the error bars represent the 95% confidence interval of the data. **(E).** Graph representing the initial Force output at 12.5 µm extension after the PSC was bound at different extensions. The bar represents the mean, and the error bars represent the standard deviation of the data.

We further investigated the higher forces generated at shorter loading extensions by measuring the initial force of PSC binding after loading at 6, 8, 10, 12, and 14 µm. The initial force is defined as the value measured when the beads were extended to 12.5 µm after loading. As expected, the most significant force, 60 pN, was observed when the PSC was loaded at 6 µm (**Figure 3E**). This dropped off at higher loading extensions (**Figure 3E**). This change likely represents a decrease in the number of contact points between the PSC and DNA. The higher forces observed at shorter loading extensions may reflect the formation of higher-order structures, creating multiple points of contact acting in parallel that can search the donor DNA with an additive force.

### Rad51 and Rad54 add stress to isolated DNA

The PSC can isolate stretches of donor dsDNA. The impact of this on the DNA within this region is unclear. Previous work has used P1 nuclease cleavage (51) and RPA binding to measure the partial melting of the DNA duplex during the homology search (29). To improve the resolution of these measurements and to understand how PSC activity impacts the remodeling of the donor DNA, we used a magnetic tweezer (MT) system to monitor the activity of Rad54 and the PSC on torsionally constrained (TC) DNA. The experimental setup consisted of a single 12.7 kbp piece of dsDNA attached to a magnetic bead on one end and to the surface of a flow chamber at the other end. Both strands are connected to the bead and the surface, making the DNA torsionally constrained (61) (**Supplemental Figure 1A**). The magnetic tweezer setup allows precise control and measurement of DNA remodeling.

For each experiment, the DNA is initially rotated through turns in the positive or negative direction, generating a hat curve based on the height of the bead (**Supplemental Figure 1A**). The curve is the result of changes in the bead height as turns are added to the DNA backbone. On either side of the hat curve, the DNA undergoes a buckling transition, which lowers the height of the bead. This transition results from a plectoneme formation at low force and is the response to torsional stress (**Supplemental Figure 1A**)(62,63). The binding and activity of Rad54 can then cause changes in the height of the bead. Events that can cause bead height changes include stretching of the regular B-form helix or the addition of isolated turns to the DNA.

The chirality of DNA remodeling can be measured by observing changes in the direction of the bead height. For example, if activity is measured at +30 magnet turns, shortening of DNA extension indicates that Rad54 has underwound the DNA. In contrast, at –60 magnet turns, underwinding the DNA will add positive turns to the plectoneme, causing the extension to increase (**Supplemental Figure 1AB**). Rad54 alone was able to shorten the height of the bead at +30 and lengthen it at –60, consistent with the addition of negative turns to the DNA (**Supplemental Figure 1B**). These data qualitatively suggest that Rad54 promotes underwinding of the DNA.

To quantitatively evaluate the number of DNA base pairs that are added to an isolated loop or remodeled during activity, Rad54 would have to tightly couple movement to the addition of turns to the DNA at 1:1. One method for evaluating this is to visually inspect activity traces to determine if the height of the bead drops below the initial extension value at –60 turns or above the initial value at +30 turns (**Supplemental Figure 1B**). Our inspection of the traces revealed a significant number of events that dropped below or above the baseline, suggesting that the motor loosely couples movement to twist. Therefore, the quantitative evaluation of Rad54 and PSC activity is limited to changes in bead extension, rather than being converted to turns or base pairs of DNA.

To measure protein activity, we analyzed the extension by taking the mean within a 5-second sliding window. The mean extension presented does not equal the total change in extension observed for the whole molecule, or whole activity trace, and successive windows could be additive or subtractive. However, our analysis provided a snapshot of local DNA remodeling events during the homology search. We measured the rate of extension using the same method, defining DNA that can be extended per second. Again, these values can be positive or negative. Here we are only reporting the positive values, as there was no significant difference between the two rates.

We measured the changes in extension of the DNA, initially starting at –60 turns, for 25, 125, and 500 pM Rad54. There was an increase in the size of extension events with increasing concentration of Rad54. The increase was not observed when ATP was omitted from the reaction (**Supplemental Figure 1C**). A mean change of 63 nm of extension was added at the highest concentration of Rad54 tested. Surprisingly, a slight change in extension occurred even in the absence of ATP, which likely stems from Rad54 binding (**Supplemental Figure 1C**). Alternatively, these small changes could also reflect noise from the measurement and represent a natural baseline. A maximum mean of 29 nm per second was observed at the highest concentration of Rad54 tested (**Supplemental Figure 1D**). Again, there was a small change in the absence of ATP, but there was no increase in activity with increasing protein concentration (**Supplemental Figure 1D**).

The lifetime of each bead extension event was measured by recording the time elapsed for events that lasted longer than 2.5 seconds and maintaining an extension at least 3 standard deviations from the baseline. The individual data points were then fit to an exponential decay curve to determine the half-life in seconds (**Supplemental Figure 1D**). There was no difference in the half-life of the events with increasing Rad54 concentrations or in the absence of ATP (**Supplemental Figure 1E**). These data indicate that Rad54 can processively relax the helical axis of the DNA in the presence of ATP.

The activity of PSCs composed of Rad51, 90-mer ssDNA, and Rad54 was measured at an initial value of –60 turns (**Figure 4AB**), at two different concentrations (**Supplemental Figure 2A**). Concentration-dependent differences were measured by keeping the ratio of Rad54:Rad51:90-mer ssDNA constant but adjusting the total concentration. For example, 500 pM Rad54 with 5 nM Rad51 and 125 pM Rad54 with 1.25 nM Rad51. These measurements were complicated by the fact that 85-90% of molecules tested would compact to the surface of the flow cell. However, this is to be expected as experiments at 0.5 pN of Force primarily resulted in compaction of the DNA via optical trapping (**Figure 1C**). For activity measurements, we focused on molecules that did not compact to the surface. We chose these molecules because they are the most likely to define single remodeling events associated with an isolated PSC. Our selection criteria were based on analyzing data until the bead extension dropped below the initial extension. At this point, we no longer used data from these molecules, even if they returned to above baseline extension.

**Figure 4:**
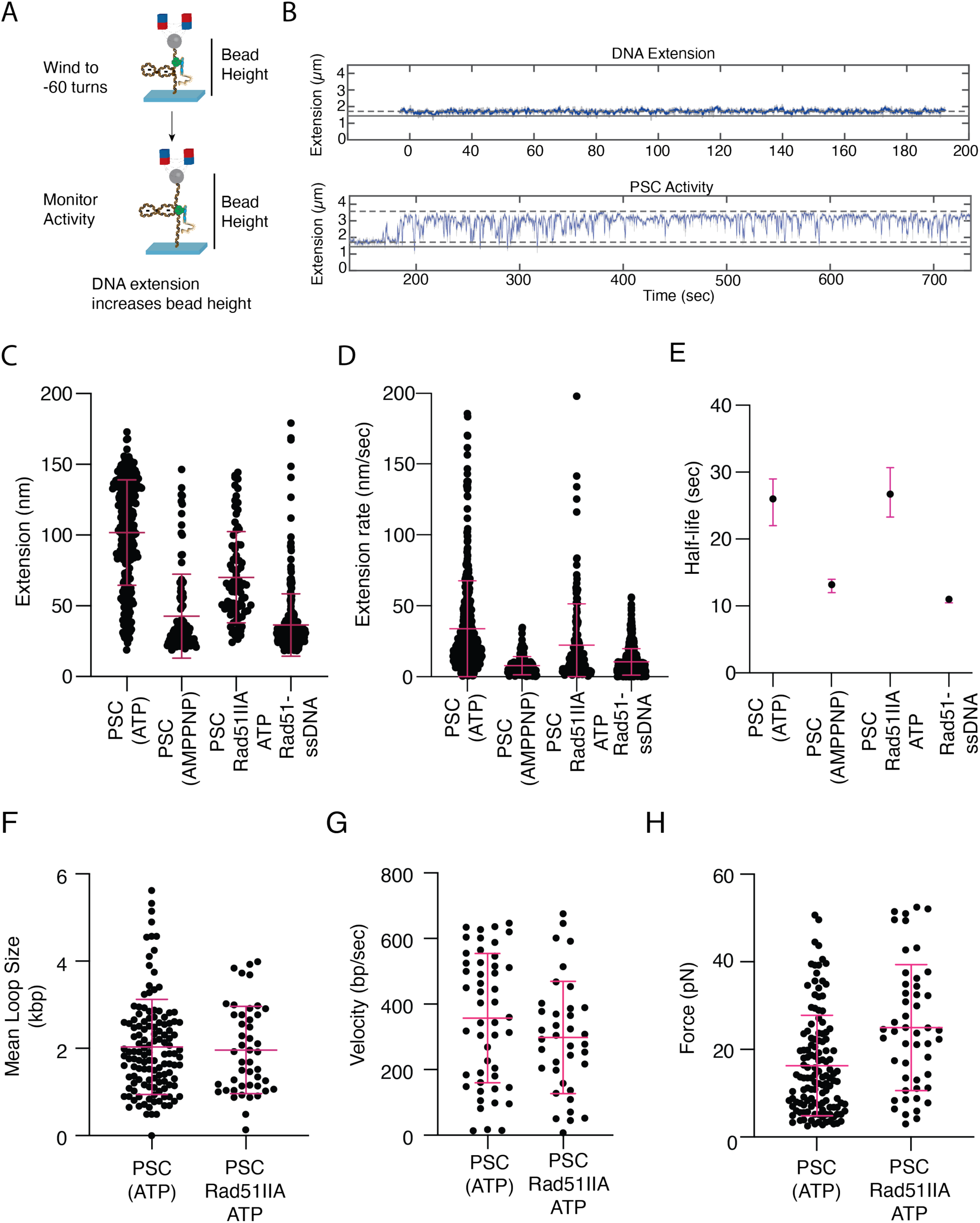
Full DNA extension requires ATP hydrolysis and Rad51-DNA binding site II. **(A).** Schematic diagram of Magnetic tweezers experiments with the PSC **(B).** Activity trace for DNA and PSC at –60 turns. The blue lines represent the change in bead height and extension of the DNA. The DNA extension was used as a baseline control to monitor deviation from DNA alone (Top). The dashed lines represent the max and min extension of the bead at –60 turns. **(C).** Dot plot representing the bead extension for PSC (ATP)(500 pM) (N=277), PSC (AMPPNP)(500 pM) (N=105), PSC with Rad51-IIA(500 pM) (N=82), and Rad51-ssDNA(5 nM) (N=309). The cross bar represents the mean of the data, and the error bars represent the standard deviation. **(D).** Graph representing the extension/second for PSC (ATP) (500 pM) (N=443), PSC (AMPPNP) (500 pM) (N=144), PSC with Rad51-IIA (500 pM) (N=161), and Rad51-ssDNA (5 nM) (N=580). The bar represents the mean, and the error bars standard deviation. **(E).** Graph representing the half-life measurements for PSC (ATP) (500 pM) (N=277), PSC (AMPPNP) (500 pM) (N=105), PSC with Rad51-IIA (500 pM)(N=97), and Rad51-ssDNA (5 nM) (N=309). The dot represents the half-life, and the error bars represent the 95% confidence interval of the fit. **(F).** Graph representing mean loop size after compaction by the PSC (N=130) and PSC with Rad51IIA (N=46). The bar represents the mean, and the error bars represent the standard deviation of the data. The PSC data is reproduced from Figure 2G **(G).** Graph representing the translocation velocity at 5 pN for PSC (N=49) and PSC with Rad51IIA (N=37). **(H).** A graph representing the force required to break contacts between the PSC and DNA for PSC (N=130) and PSC with Rad51-IIA (N=49). The bar represents the mean, and the error bars represent the standard deviation of the data. As in Figure 2, the mean force is a measure of intermediate peaks in the re-extension curves.

We observed a concentration-dependent increase in the change in extension observed between 500 pM and 125 pM PSC (101 versus 79 nm, p <0.0001) (**Supplemental Figure 2B**). There was also an increase in the half-life of events formed at higher PSC concentrations (∼10 sec longer) (**Supplemental Figure 2C**). However, there was no difference observed in the extension per second, suggesting the rate was unaffected by concentration (**Supplemental Figure 2D**). From these measurements, we conclude that the stability and size of remodeled DNA were affected by the concentration of the PSC. These differences could reflect differences between 1D translocation-based search versus a 3D diffusion-based search, as PSCs with longer lifetimes are more likely to move along the DNA processively.

By performing activity measurements at –60 turns, we were able to measure the interaction of Rad51-ssDNA filaments with the donor DNA without Rad54. Interestingly, Rad51-ssDNA was able to extend DNA by around 36 nm. This was 70 nm smaller than the corresponding PSC measurements (**Figure 4C**). Additionally, the change in extension per second and the half-life of the events were significantly shorter than the PSC (**Figure 4DE**). Interestingly, the change in extension per second was ∼10.4 nm/sec, which, based on the structure of Rad51-ssDNA filaments, is the helical pitch. This would correspond to roughly one turn of a Rad51 helix or 6.4 protomers and roughly 18 bp. This is only slightly longer than the 12 bp and 4 protomers observed for RecA filaments during probing of donor DNA during the homology search (20). These differences are likely within the error of the measurement. We also evaluated the contribution of ATP hydrolysis by forming PSCs with AMPPNP instead of ATP. Under these conditions, the extension, extension/second, and half-life of the events were all the same as Rad51-ssDNA alone (**Figure 4CDE**). These data suggest that ATP hydrolysis by Rad54 is required for the additional DNA remodeling and stabilization of the PSC on the DNA.

To better understand the independent contributions of Rad54 and Rad51 in this experiment, we used a mutant form of Rad51 with three amino acid substitutions: R188A, K361A, and K371A. These substitutions disrupt the DNA binding site II within Rad51 and can be referred to as Rad51IIA (19). This site is conserved in all recombinases, and mutations in these residues fail in the homology search (7,32) and strand exchange (19). We reasoned that Rad51IIA would fail to bind the donor dsDNA, and PSCs with Rad51IIA would have smaller extensions, extension rates, or shorter lifetimes. At 500 pM PSC with Rad51IIA, there was a loss of around 31 nm from the extension events, comparable to the length of the Rad51-ssDNA filaments (**Figure 4C and Supplemental Figure 2B**). At lower concentrations, this loss was less (∼20 nm) (**Supplemental Figure 2B**). However, this could be due to the reduced DNA extension by Rad54 under these conditions. There was also a difference observed between the PSC and PSC with Rad51IIA in the extension per second (**Figure 4D and Supplemental Figure 2D**). A difference of around 10 nm per second, or the equivalent of Rad51 alone, and a single helical turn of the recombinase filament. There was no observed difference in the half-life between PSCs and PSCs with Rad51IIA. The absence of Rad51-ssDNA extension indicates that the filaments are no longer able to interact with the donor DNA, and we conclude that Rad54 can increase the binding and stability of Rad51 filaments in the absence of sequence homology by underwinding the DNA.

We asked whether Rad51 contributed to the formation of the larger loops observed via optical trapping, or if it simply remodeled DNA in an isolated region. If Rad51 contributed to the formation of larger loops, then the Rad51IIA mutant would show a reduction in loop size or loop extrusion rate. This was not the case, and we observed that the mean loop size was comparable to WT at 0.5 pN force (**Figure 4F**). We also did not observe a reduction in the translocation rate for PSCs with Rad51IIA at 5 pN force (**Figure 4G**). There was a small increase in the force required to separate intermediate contacts that formed during compaction upon re-extension (**Figure 4H**). From this, we conclude that there is no significant impact of Rad51-ssDNA binding on the formation of large, isolated loops. Instead, it suggests that Rad51 binding primarily affects the extension of the donor DNA.

We measured the activity of the PSC and Rad51-ssDNA at +30 turns. Under these conditions, the DNA is overwound, reducing the affinity of Rad51-ssDNA for the donor DNA. For Rad51-ssDNA alone, there were only 10 active molecules, which did not add significant changes to the DNA. In contrast, the PSC extended around 70 nm, illustrating significantly greater activity (**Supplemental Figure 3A**). Furthermore, these events had longer half-lives than Rad51-ssDNA alone (**Supplemental Figure 3B**). These data suggest that Rad54 can significantly aid Rad51 activity on overwound DNA.

At the end of each experiment, the magnet was rotated from –70 to 70 and back. This could identify any changes to the hat curve that occurred during the reaction. For Rad51-ssDNA alone, a translational shift in the hat curve was observed, similar to that observed for RecA-ssDNA in the presence of AMPPNP (**Supplemental Figure 4A**) (26). Interestingly, when the same measurement was made with the full PSC, the negative side of the hat curve became flatter and did not undergo a buckling transition (**Supplemental Figure 4B**). Similar observations were made for the PSC with Rad51IIA (**Supplemental Figure 4C**), with the exception that these hat curves were shorter. The slope of the hat curve from –70 to –20 turns was used to make a comparative assessment. The value was substantially different between Rad51-ssDNA, the PSC, and PSC with Rad51IIA (**Supplemental Figure 4D**). These data are consistent with donor DNA that is prone to melt instead of buckle, and this effect was dependent on Rad54 concentration in the PSC (**Supplemental Figure 4D**). Rad54 alone did not cause this outcome (**Supplemental Figure 4D**). This suggests that DNA bound by a PSC is predisposed to melt upon the addition of superhelical stress or torsion. In this scenario, as additional turns are added to the DNA, the DNA will go through a melting transition and not form a plectoneme. One interpretation is that Rad54 predisposes the donor DNA to melt when complete homology is recognized by Rad51, representing a novel mechanism by which D-loops might be stabilized.

### Mutants defective in DNA compaction fail to form D-loops *in vivo*

We have previously characterized mutations in Rad54 that result in defects in processive movement of the PSC along donor DNA. These mutant versions of Rad54 are defective for homology search *in vitro* (30). Defects stem from insufficient ATP hydrolysis required for translocation. We reasoned that these Rad54 mutants (Rad54 R272Q and Rad54 R272A) would be defective in generating loops, extending the DNA, or both. We initially measured the activity of PSCs with these mutants by optical trapping experiments (**Figure 5AB**). As expected, there was a 3.2. fold reduction in the loop extrusion rate for the PSC with Rad54R272Q and a 5.1-fold reduction for the Rad54 R272A (**Figure 5C**). The mutant forms of Rad54 also exhibited a ∼800 bp reduction in loop size (**Figure 5D**). Re-extension of the DNA after compaction required 1.75 times the mean force to disrupt intermediate contacts for Rad54 R272Q and 2.3 times more force to disrupt intermediate contacts. From this, we conclude that PSCs with Rad54 mutants are defective in loop compaction, which is likely due to an increased affinity for dsDNA, reducing processive movement.

**Figure 5:**
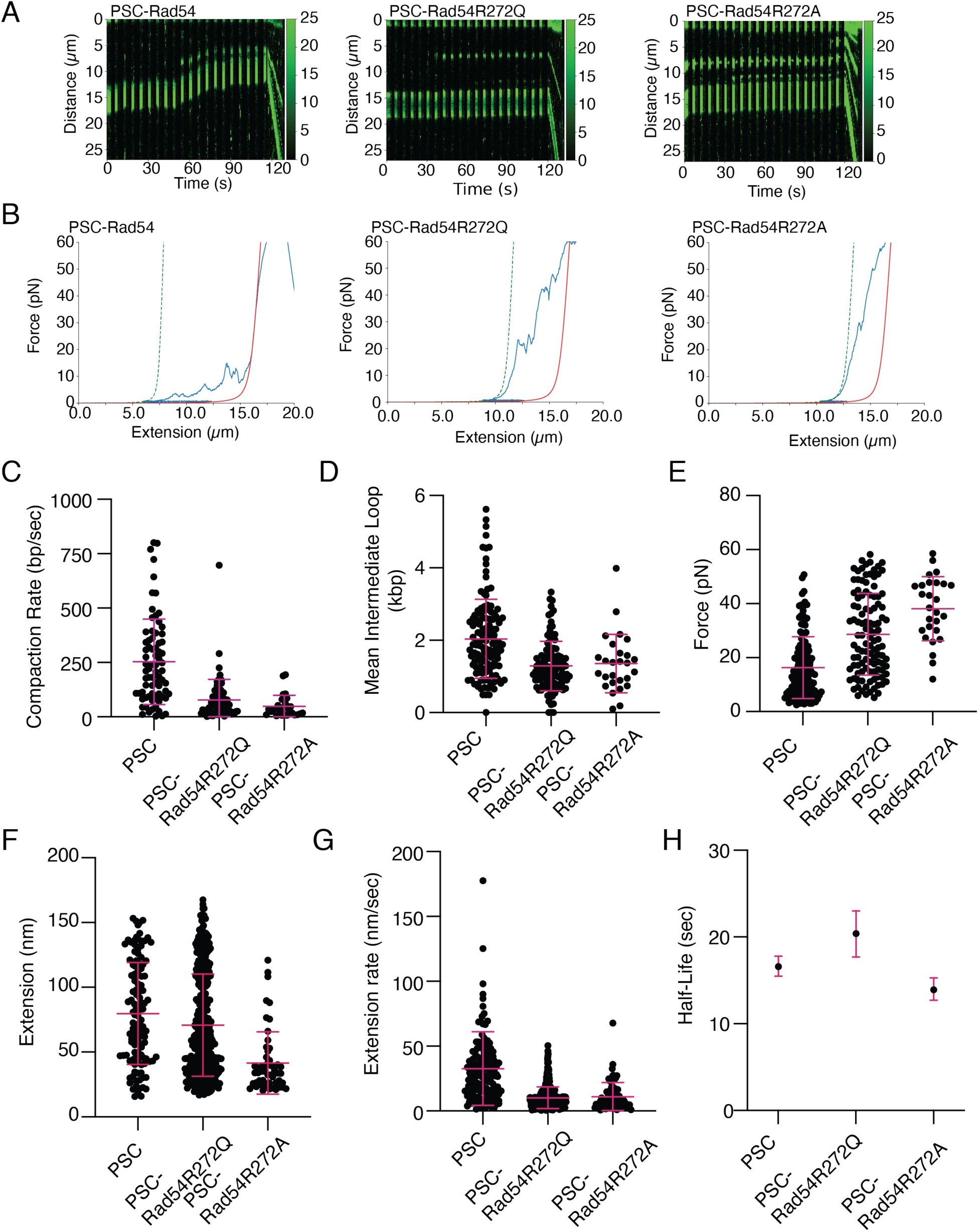
Defects in Processive translocation alter loop dynamics. **(A).** Representative kymographs for PSC, PSC ( Left) with Rad54R272Q (Middle), and PSC with Rad54R272A (Right). **(B).** Representative post-compaction FE curves for PSC (Left), PSC with Rad54R272Q (Middle), and PSC with Rad54R272A (Right). The blue line represents the measured forces, and the red line represents the theoretical force extension curve for the DNA. The green dashed line represents the initial point of deviation from the theoretical FE curve **(C).** Dot plot comparing the loop extrusion rate for PSC (N=74), PSC with Rad54R272Q (N=78), and PSC with Rad54R272A (N=28) at 0.5 pN. The bar crossbar represents the mean of the data, and the error bars represent the standard deviation. The data for the PSC is reproduced from Figure 1D. **(D).** A dot plot represents the mean intermediate loop size for PSC (N=130), PSC with Rad54R272Q (N=111), and PSC with Rad54R272Q (N=27). The crossbar represents the mean of the data, and the error bars represent the standard deviation. The data for the PSC is reproduced from Figure 4. **(E).** A dot plot represents the force required to disrupt interactions in compacted DNA structures for PSC (N=130), PSC with Rad54R272Q (N=111), and PSC with Rad54R272Q (N=27). The crossbar represents the mean of the data, and the error bars represent the standard deviation. The data for the PSC is reproduced from Figure 4. **(F).** Graph representing extension for the MT data of PSC (125 pM) (N=120), PSC with Rad54R272Q (N=362), and PSC with Rad54R272A (N=54). The crossbar represents the mean, and the error bars represent the standard deviation. The PSC data is reproduced from Supplemental Figure 3. **(G).** Graph representing the extension per second for the PSC (125 pM)(N=148), PSC with Rad54R272Q (125 pM) (N=402), and PSC with Rad54R272A (125 pM) (N=70). The line represents the mean, and the error bars represent the standard deviation of the experiment **(H).** Graph representing the extension half-life for PSC, PSC (N=277) with Rad54R272Q (N=362), and PSC with Rad54R272A (N=54). The dot represents the half-life and the error bars the 95% confidence of the fit.

We next evaluated how these substitutions affected the change in DNA extension. These experiments were performed at 125 pM PSC. Under these conditions, the WT PSC extended the DNA around 80 nm (**Figure 5F and Supplemental Figure 5ABC**). The PSC-Rad54R272Q mutant extended DNA around 70 nm, or around 88% the WT value. The PSC Rad54R272A extended the DNA 41 nm, representing a 2-fold reduction (**Figure 5F**). There was a significant reduction in the DNA extended per second, with the Rad54 R272Q and the Rad54 R272A mutant both having a 3.2-fold defect (**Figure 5G**). The extension half-life was slightly longer than the WT for the Rad54 R272Q, but there was no difference for the Rad54 R272A, suggesting the half-life of these events was comparable to WT (**Figure 5H**). While these mutations are defective in all aspects of the PSC activity, the most severe defects are associated with DNA compaction and the rate of DNA extension (30). Together, these data suggest that the ability to act processively is critical for PSC function.

To determine if these mutants were defective in target search *in vivo,* we utilized a reporter assay from *S. cerevisiae* that measures the amount of D-loop capture during early strand exchange (64,65). The experiment is based on an HO nuclease site located at a specific location on chromosome V, with a homologous region of DNA located on chromosome II (**Figure 6A**). The HO nuclease can be induced with galactose, and the broken recipient DNA can undergo a homology search and D-loop formation. Nascent D-loops can be detected by using psoralen cross-linking to trap the structure in cells. Psoralen is a DNA intercalating agent activated by UV whose activity depends on DNA topology. We monitored nascent D-loop capture at 3 hrs and found that in WT strains, D-loops were captured at an efficiency of 3.3 ×10 +/−0.06×10^−2^ (**Figure 6B**). In contrast, the *rad54* had a capture efficiency of 1.7×10 +/−0.096×10^−4^ (**Figure 6B**), representing a 200-fold reduction from WT. The *rad54R272Q* and *rad54R272A* strains both captured D-loops 80-fold less efficiently than WT (**Figure 6B**). This was surprising because these mutations are less severe than those in *rad54* in methyl methanesulfonate (MMS) sensitivity assays, and *in vitro*, longer fragments of ssDNA can form D-loops (30). We expected both mutants to retain partial function. However, this was not the case. These experiments demonstrated that substitutions that cause defects in PSC processivity and the addition of DNA stress result in an 80-fold defect in D-loop capture and are defective for homology search *in vivo*.

**Figure 6:**
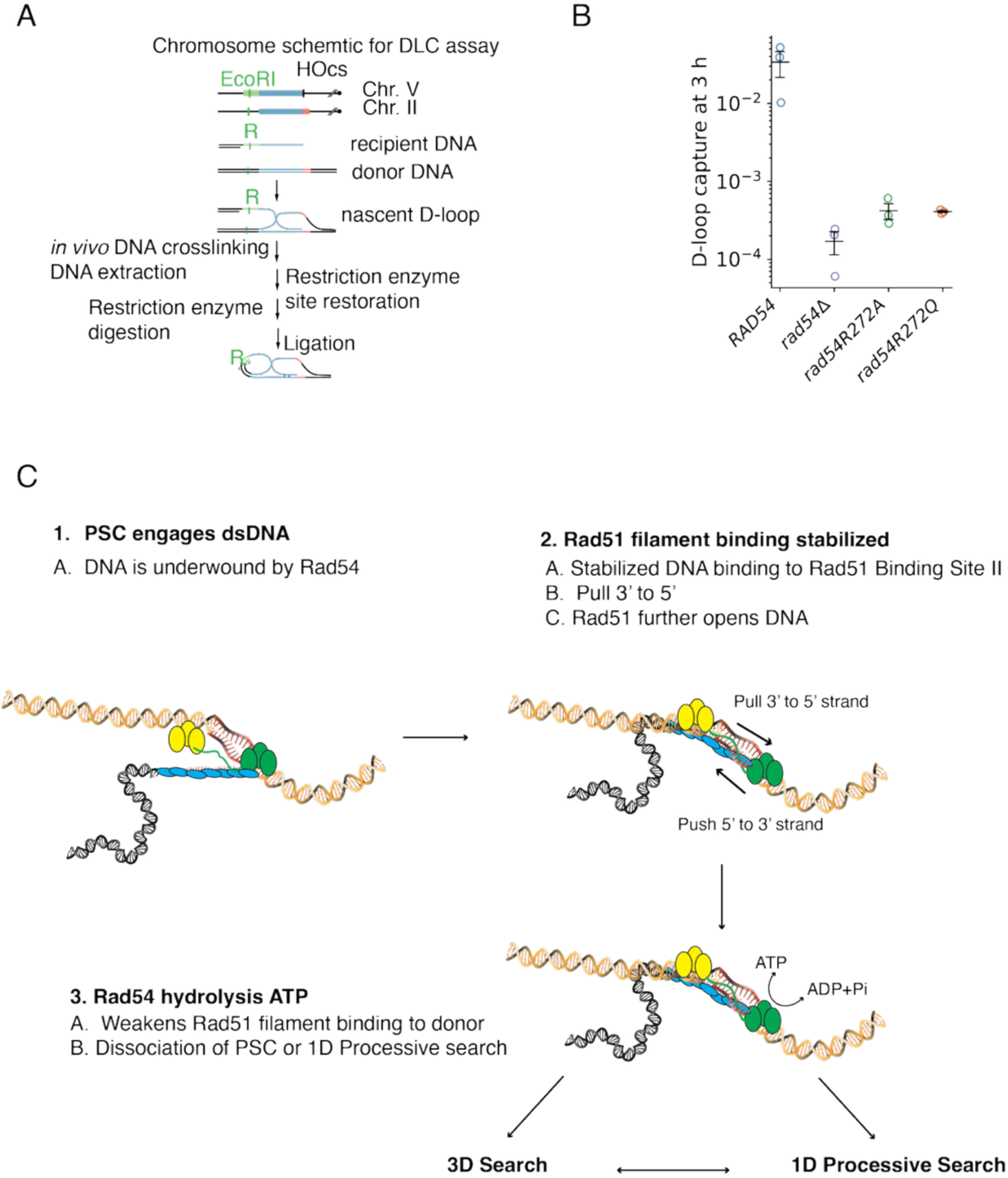
Failure in processive search limits D-loop formation *in vivo*. **(A).** Cartoon diagram illustrating the D-loop capture experiment used in *S. cerevisiae* cells to determine the efficiency of nascent D-loop capture in cells. **(B).** Dot plot representing the nascent D-loop capture efficiency for *RAD54, rad54, rad54R272A*, and *rad54R272Q.* The crossbar represents the mean of the data, and the error bars represent the standard error of at least three independent experiments. **(C).** Cartoon diagram illustrating the model for cooperative homology search by Rad51 and Rad54.

## Discussion

The RecA family of recombinases forms filaments that can introduce both linear (tension) and rotational (torsion) forces onto duplex donor DNA. However, to introduce these forces during the homology search and strand exchange, they must efficiently bind to the donor DNA. Both Rad51 DNA-binding sites prefer single-stranded DNA (ssDNA), and relaxing the superhelical structure of the donor DNA substrate is an efficient way to enhance the binding of Rad51-ssDNA filaments to the donor duplex prior to the recognition of homology. Here, we demonstrated that the ATP-dependent motor protein Rad54 can catalyze Rad51-ssDNA binding in the absence of homology by providing assisting forces. Both in the form of linear (tension) and rotational (torsion) stress placed on DNA, with the latter requiring the isolation of a donor DNA loop by Rad54. Importantly, hydrolysis of ATP can remove the forces placed on DNA, temporarily reducing the affinity of Rad51-ssDNA filaments for the donor DNA. The ability of Rad54 to regulate Rad51-ssDNA binding assists the search for homologous bases and is essential for sequence identification and D-loop formation *in vivo*.

### The activity of the PSC donor DNA structure

DNA loops are common in biology and generally depend on the length of DNA between two points of protein-DNA contact (58,66,67). When an active motor protein captures a loop, additional DNA can be added to the loop through a process known as loop extrusion. Loop extrusion can modify the topology within the loop as long as the turns added to the DNA remain constrained. The PSC can form short, isolated loops on the DNA that are ∼2-2.5 kbp in size. Our data suggests that the PSC regulates the superhelical density of these loops.

The donor DNA can be bridged by Rad54, a proposal that has been made before (68). DNA bridging can exist in either a cis-or trans-configuration, creating local domains that are underwound and can be probed by Rad51. The twin domain supercoiling model suggests that if regions of underwound DNA are present, then there must also be regions of overwound DNA to accommodate the negative supercoiling. When Rad54 contacts occur in cis, overwound DNA may exist between these two points of contact, and these regions may be inaccessible to Rad51. However, as the molecules move along the DNA through translocation or compaction, the underwound and overwound regions may shift. Additionally, as the PSC dissociates to explore new areas of the genome, topologically isolated domains will form and dissolve to accommodate the search. In a cellular context, the PSC is likely built on a longer piece of resected ssDNA with multiple short Rad51 filaments. The ability of each of these regions to probe multiple pieces of donor DNA simultaneously may significantly shorten the amount of time required to find a homologous sequence (**Supplemental Figure 6**).

### Relaxing the superhelical axis of DNA

In many molecules we analyzed via magnetic tweezers, the donor DNA was compacted to the surface. This is likely due to higher-order protein-DNA interactions and is like the compaction observed via optical trapping. Most of our analysis focused on measurements that did not completely reach the surface, which likely represent individual units that were measured to be roughly 2 kbp in length. Overall, both Rad51 and Rad54 could act to add negative turns to the DNA. We were able to differentiate the contributions of Rad54 and Rad51 by using the Rad51IIA mutant, which disrupts the DNA binding site II on the recombinase filament. This did not result in a reduction of bead extension lifetime but did result in a reduction in DNA extension rate comparable to the length of a single turn of a Rad51-ssDNA filament. This is consistent with a loss of Rad51-ssDNA binding to the DNA. The stability of binding in the context of the PSC was dependent on Rad54 activity. Experiments at +30 turns, which overwind the DNA, did not produce extended binding events in the absence of Rad54. However, Rad54 was able to promote the binding of Rad51 filaments in overwound regions. Highlighting the role of Rad54 in modulating local DNA structure. This function may ultimately be important for replication restart (69) or dealing with parts of the genome that are positively supercoiled.

Our general model for the remodeling of donor DNA is that Rad54 can promote both linear (tension) and rotational (torsion) stress on the DNA, resulting in underwound donor DNA. This enhances the affinity of Rad51-ssDNA filaments for DNA, allowing them to associate with and sample the donor DNA sequence (**Figure 6C**). Importantly, the stress placed on DNA can be lost when Rad54 hydrolyzes ATP, causing the affinity of Rad51-ssDNA filaments to go down. This action could occur if tension is lost on the donor DNA, or positive turns are added to the loop from a trailing Rad54 motor (**Figure 6C).** The result of ATP hydrolysis is 1D translocation or dissociation and 3D diffusion to a new target. Unless the filament is stabilized by homology recognition, which will result in strand exchange.

### Failures in homology search

Mutations that disrupted ATP hydrolysis by Rad54 also failed to promote homology search and nascent D-loop capture in *S. cerevisiae*. *In vitro,* these defects reflect the increased stability of looped intermediates, resulting in decreased loop formation rates and smaller loop sizes. These outcomes are reflective of an enzyme that fails to turn over, resulting in a homology search complex that becomes stuck to the donor DNA. The outcomes of these experiments underscore the importance of Rad54 turnover in the homology search process, as it facilitates 1D translocation and regulates PSC stability during 3D search.

Recently, the participation of Rad54 in the homology search was measured in live cells. In baker’s yeast, the deletion of Rad54 leads to the lengthening of Rad51 filaments (70). Possibly due to the loss of donor DNA compaction, resulting in an apparent increase in filament size. In human cells, RAD54L was required to resolve RAD51 foci during the homology search. However, this mechanism is unclear. A cooperative interaction between human RAD54L and Cohesin was also observed during the homology search (71), and could occur by regulating the forces placed on the DNA. Further work will be needed to understand the direct biochemical relationship between the SMC protein Cohesin and the PSC.

It remains to be seen if RAD54L will make similar contributions during the homology search in human cells. Human RAD51 is more effective at binding to dsDNA in the context of the PSC, and additional factors, such as BRCA2 and RAD51AP, have evolved to regulate RAD51 binding to DNA (72–76). While RAD54L is required for the resolution of RAD51 foci in human cells, this could be due to the removal of RAD51 filaments or modulation of PSC binding by donor DNA remodeling. Resolution of these possibilities will require the use of hypomorphic alleles specifically designed to separate these functions.

### Limitations of our study

A limitation of our study is that it is unclear how well conserved this mechanism might be between yeast and human Rad54. Although the proteins are 50% identical, there could be differences in overall function dependent on context. Additionally, our experiments don’t reveal the actual outcomes of sequence recognition and the formation of full D-loops. Future work will be needed to address these two key points.

### Data sharing plan

Data, including kymographs, will be submitted to the Medley repository upon acceptance of this manuscript. All analysis codes are uploaded to GitHub. Additionally, all other data will be available on request.

## Acknowledgments

We want to acknowledge members of the Crickard and Wang laboratory for helpful discussions on this manuscript. This work is supported by National Institutes of Health grant 5R35GM142457 (to J.B.C) M.D.W is an Howard Hughes Medical Investigator.

## Author contributions

MVW carried out all single-molecule experiments, purified proteins, analyzed the data, helped with figure preparation, and contributed to the writing of the manuscript. JH performed the *in vivo* D-loop capture assay and helped with figure preparations. MW generated templates for MT experiments and assisted with single-molecule experimental design. JQ assisted with MT experiments and MT experimental design. JI provided experimental support with MT and C-trap experiments and helped with data analysis. MDW provided instruments and resources and proposed single-molecule experimental approaches. JBC provided resources, proposed experimental approaches, guided experimental direction, and wrote the manuscript with input from all authors.

## Declaration of interests

The authors declare there are no competing interests.

## Declaration of generative AI and AI-assistance

Grammarly AI assisted editing was used in the preparation of this manuscript. No generative AI was used.

## Materials and Methods

### Protein purification

Rad54, Rad54R272Q, Rad54R272A, Rad51 and Rad51-IIA were purified as previously described (29). In brief, a protease deficient yeast strain was transformed with GFP-GST-Rad54, GFP-GST-Rad54 R272Q or GFP-GST-Rad54 R272A, on 2–micron plasmids under the control of the Gal1/10 promoter. Cells were grown in Yeast Nitrogen base (–URA) plus 3% Glycerol and 2% lactic acid. When the cells reached an OD of 1.5, expression was induced by adding 2% galactose for 6 hours. Cells were harvested and stored at–80°C.

Cell pellets were resuspended in Rad54 resuspension buffer (30 mM Tris–HCl [pH 7.5], 1 M NaCl, 1 mM EDTA, 10% glycerol, 10 mM BME (β-mercaptoethanol), protease inhibitor cocktail (Roche Cat. No. 05892953001), and 2 mM PMSF. Cells were disrupted by manual bead beating, and the lysate was clarified by centrifugation at 26,500xg for 1 hour. The lysate was fractionated by ammonium sulfate (AS) precipitation. AS was gradually added with mixing to a final concentration of 20% followed by centrifugation at 10,000 x g for 10 minutes. The supernatant was discarded, and the AS concentration was raised to 50% followed by centrifugation at 10,000 x g for 10 min. The protein pellet was resuspended in PBS (phosphate buffered saline) plus 1M NaCl and 10 mM BME. The resulting re-suspended protein was then bound to pre-equilibrated GST resin in batch for 1 hour at 4°C. The GST resin was washed 2x with PBS plus 1000 mM NaCl, and 2x with PBS plus 500 mM NaCl. The protein was eluted in 20 mM glutathione in PBS plus 500 mM NaCl. The peak fractions were pooled and then applied to a Sephacryl S–300 High Resolution gel filtration column (GE Healthcare, Cat. No. 17–0599–10) pre–equilibrated with Rad54 SEC buffer (30 mM Tris–HCl [pH 7.5], 500 mM NaCl, 1 mM EDTA, 10% glycerol, and 10 mM BME. The peak was pooled and dialyzed against Rad54 SEC buffer plus 50% glycerol and stored at –80°C in single-use aliquots.

6xHis–SUMO–Rad51 or 6xHis-SUMO-Rad51IIA was transformed into *E. coli* BL21 (DE3) Rosetta2 cells and grown to an OD_600_ of 0.4–0.6 at 37°C to. Expression was induced by addition of 0.5 mM IPTG for 3 hours at 37°C Cells were harvested and stored at –80°C. Cells were lysed by freeze–thaw in Cell Lysis Buffer (CLB:v30 mM Tris–HCl [pH 8.0], 1 M NaCl, 10% glycerol, 10 mM imidazole, 5 mM BME, and protease inhibitor cocktail (Roche Cat. No. 05892953001)). Crude lysates were sonicated for 6 pulses of 30 seconds on and 2 minutes off, then clarified by centrifugation at 26,500 x g. The extract was precipitated with 50% ammonium sulfate and centrifuged at 26,500 x g for 10 minutes. The Pellet was resuspended in CLB and bound to 1 mL of pre–equilibrated Ni–NTA resin for 1 hour with rotation at 4°C. The resin was washed 3X with CLB and eluted in CLB+200 mM imidazole. The protein was mixed with 400 units of the SUMO protease Ulp1 and dialyzed overnight at 4°C into Rad51 buffer (30 mM Tris–HCl [pH 8.0], 150 mM NaCl, 1 mM EDTA, 10% Glycerol, 10 mM imidazole). The 6xHis–SUMO tag and SUMO protease were removed by passing the dialyzed proteins over a second 1 mL Ni–NTA column. The purified Rad51 was then stored at –80°C in single use aliquots.

### DNA Template Construction

The torsionally constrained DNA template was generated by adding a ∼500-bp multi-labeled adapter at each end with a center segment for a total length of 12,688 bp DNA center (77). The center segment was PCR amplified from λ-DNA (NEB, N3011S), then double digested with AvaI (NEB, R0152S) and BssSI-v2 (NEB, R0680S) to produce the unique overhangs for ligation. To make the 500-bp multi-biotin-labeled and multidigoxigenin-labeled adapters, we performed PCR amplification from plasmid pNFRTC (pMDW111) with either 24% of dATP replaced by biotin-14-dATP or 24% of dTTP replaced by digoxigenin-11-dUTP, followed by restriction enzyme digestion with BssSI and AvaI, respectively. The ∼500-bp multi-labeled adapters with unique overhangs were ligated to the 12,688 bp center segment. The lambda template ligation product was gel-purified, and aliquots were stored at –20 °C.

### Magnetic Tweezers

Experiments on the magnetic tweezer (MT) were performed on a custom-built instrument (61), allowing for bulk analysis of multiple DNA tethers under a constant force. In each chamber, between 30-50 tethers remained constrained (TC) throughout the experiment and were affected by the rotation of the magnetic beads. Chambers for the MT were prepared by nitrocellulose coating (1-2% collodion in Amyl acetate) on two microscope coverslips, forming a flow cell with the nitrocellulose surfaces facing inward. The surface of the chamber was then functionalized with anti-digoxygenin (Vector Labs MB-7000), passivated with 1.25 mg/mL β-Casein (Sigma C6905), incubated with 1-2 pM 12.7 kb λ-DNA template, and lastly incubated with streptavidin-coated paramagnetic beads (Invitrogen 65601). To ensure no free beads remained in the solution, a buffer exchange was done to start all chambers in HR Buffer (30 mM Tris-OAc [pH 7.5], 10 mM Mg(OAc)_2_, 50 mM NaCl, 1.5 mg/mL β-Casein, 1 mM DTT). Before introducing Rad54 to the sample chamber, DNA tethers were assayed to determine torsional constraint and to provide a baseline for unadulterated tether behavior. The tethers were twisted via rotation of a magnet until overwound and underwound, leading to the generation of a buckling curve. Tethers were also left in an overwound or underwound state for 2 minutes compared to tethers with bound protein. To load protein, the force in the chamber was increased to 6 pN while protein was flown in for 2 minutes, after which the tethers were over/underwound, and the force dropped to the experimental force (typically 0.5 pN). A monitor was engaged for 10 minutes, after which the initial buckling curve (hat) was regenerated for further analysis. Changes to the hat curve were analyzed by measuring the slope of the negative side of the curve. This was compared to DNA without proteins.

### MT Data Analysis

Activity traces were collected over ten-minute periods. Active traces were determined by using the fluctuation of the torsionally constrained DNA alone as a baseline. Changes in extension were identified by determining local maximum/minimum within an extension event. A change in extension was considered an event when it exceeded 3 standard deviations from the baseline, as determined from the DNA alone. Positive rates were determined by identifying the slope leading to a local maximum. Negative rates were determined by identifying the slope following a local maximum. Slopes were linear between maxima/minima. Extension data was smoothed by applying a 5-second sliding window. The mean extension for a given DNA template was determined by fitting to a Gaussian distribution. The lifetime of each extension event is determined by setting a threshold of the mean extension of naked DNA under torsion and analyzing traces that extend + 3 standard deviation (std) from the baseline. Any events that crossed this threshold lasting greater than 2.5 seconds were considered an active trace. The lifetime of events can be determined by the time between points that are greater than 3 std from the mean. These lifetimes fit an exponential decay curve. The code used for all MT analysis can be found on Github.

### Lumicks Confocal Microscopy with optical trapping

All fluorescent experiments were conducted on a LUMICKS C-Trap instrument, allowing for the combination of an optical dual-trap with confocal imaging microscopy. Excitation lasers at 488, 532, and 647 nm allowed for the excitation of GFP (Rad54), dsDNA (Sytox Orange), and ssDNA (Atto-647N), respectively. Before experiments, the five-channel laminar flow cell (Model C1) was passivated using 0.5 mg/mL B-Casein (Sigma C6905) in a standard Running Buffer (PBS, 1.5 mM sodium azide, 0.5 mM EDTA). To form DNA tethers, streptavidin-coated polystyrene beads (0.004% w/v; LUMICKS; diluted in Running Buffer, 5 µm size) were trapped and moved briefly into the biotinylated λ-DNA channel (8 pg/µL; LUMICKS; diluted in Running Buffer). After tether formation, the traps were moved to the HR Buffer in channel 3 and stretched to ensure single tether formation. All protein was loaded into channel 4 in HR Buffer + 1 mM ATP. For catching beads, forming tethers, and loading protein, the flow was kept at a constant 0.2 +/−0.05 bar. The trapping power was set to 7.5% during data collection, leading to a trap stiffness of ∼0.07 pN/nm. Kymographs were collected with 0.5 millisecond pixel dwell time for each 100 nm pixel. The kymograph frame rate alternated between 2 (0.5 ms exposure) and 10 (0.25 ms exposure) frames per second, depending on experimental needs. Unless otherwise dictated, kymographs were collected using staggered excitation lasers, where each laser would be on for 1 second and off for 2 seconds. For conditions where the 532 lasers were not used, excitation alternated between 488 and 647 lasers in 1-second increments. These were temporally offset to prevent bleed-through of the channels.

### Lumicks Confocal microscope with optical trap (C-trap) Experimental Protocols

The experimental protocol for LUMICKS experiments was separated into three automated scripts. All experiments were followed by overstretching the DNA at a constant rate.

### Force Clamp Experiments

Captured DNA tethers were moved into Channel 4 at a high (16 μm) extension. Protein was loaded via 60 seconds of flow. After protein loading, a force clamp was entered at 0.5, 1, 2, or 5 pN, and a kymograph was generated using 488/532/647 at 2 frames per second (FPS).

### Force extension measurements

Captured DNA tethers were moved into Channel 4 at varying extensions (6, 8, 10, 12, 14 μm). Protein was loaded via 30 seconds of flow. After loading protein, the tether was moved to a 12.5 μm extension, equivalent to 0.5 pN of force on naked DNA. Kymographs were generated using 488/647 lasers at 2 frames per second (FPS). Force-extension curves were also collected during the experiment.

### Lumicks Data Analysis

Raw data exported from LUMICKS Bluelake as .h5 files were processed in Spyder using Python 3.10 and custom-generated software. Kymographs were generated, and all measurements were generated from the intensity data included in these kymographs. Particle tracking was done by manually selecting bound proteins. The resulting particles were used to measure velocities, binding lifetimes, and fluorescent intensities. Translocation velocities were calculated by measuring the distance changed over unit time. Compaction measurements, the degree of compaction, and the compaction rate were analyzed by changes in the distances between the two beads as measured by the extension curve. Re-extension of the compacted DNA was performed at a defined rate, and measurements were performed by analyzing the Force extension curve. The size of defined loops was determined from the size of the DNA when a loop disruption resulted in a return to a theoretical re-extension curve. Each loop disruption was included for further analysis. Analysis of force measurements was performed by analyzing the Force-Extension curve during translocation. Max force intensities during translocation were used to generate mean force output.

## Yeast Strain Construction

The yeast strains used in this study was a kind gift from Wolf Heyer. The *rad54* strains were generated by gene knockout using a KanMX cassette. The *rad54R272Q*/A strain was generated by using gene replacement with a PCR product to form a *pRS305-rad54R272Q/A-KanMX* plasmid. WT *RAD54* was replaced similarly, generating *RAD54-KanMX*, which was used as the WT.

## DLC assay

DLC assay was performed as described (64,65). Yeast cells were grown in a 5 mL Yeast extract Peptone (YP) medium supplemented with 2% dextrose and 4% adenine sulfate overnight. The second day, the culture was diluted by 10-fold in 5 mL YP + 3% glycerol + 2% lactate + 4% adenine sulfate and grown for around 8 hours. Then the culture was inoculated into 100 mL of YP + 3% glycerol + 2% lactate + 4% adenine sulfate medium with an initial OD_600_ ≈ 0.006 and grown for 16 hours. A 5x psoralen stock solution (0.5 mg/mL trioxsalen in 200-proof ethanol) was made in a 50-mL aluminum foil-covered tube and dissolved on a shaker at room temperature overnight with gentle rocking. The next day, the culture should have an OD_600_ of 0.3-0.8. 7.5 OD_600_ of cells was collected as time 0 control, centrifuged at 2,246 x g rpm, 4 °C for 5 minutes. The cell pellets were resuspended in 1x psoralen buffer. The 1x psoralen buffer was prepared by diluting 5x psoralen in 200-proof ethanol before collecting cells. The resuspended cells were plated in a 60 mm petri dish, put 2-3 cm below a UV light source with the lip removed atop a pre-chilled metal block. The cell samples were exposed under the UV light for 10 minutes with gentle shaking to crosslink DNA. The cells were transferred to a 15-mL Falcon tube. The petri dish was rinsed with TE1 solution (50 mM Tris-Cl pH 8.0, 50 mM EDTA pH 8.0) and the TE1 buffer was poured together with cells. The cells were then centrifuged at 2,246 x g, 4 °C for 5 minutes again. The pellets were saved at –20 °C. Galactose was added into the culture to a final concentration of 2% to induce DSBs. Cells were collected at designated time points as described above.

The cell pellets were thawed on ice, then resuspended in spheroplasting buffer (0.4 M sorbitol, 0.4 M KCl, 40 mM sodium phosphate buffer pH 7.2, 0.5 mM MgCl_2_) and transferred to a 1.7 mL microfuge tube. The cells were spheroplasted in zymolyase solution (2% glucose, 50 mM Tris-Cl pH 7.5, 5 mg/mL zymolyase 100T) at 30 °C for 20 minutes. The cells were washed by spheroplasting buffer for three times at 2500 xg and restriction enzyme buffer (RE buffer, 50 mM potassium acetate, 20 mM Tris-acetate, 10 mM magnesium acetate, 1 mg/mL BSA) at 16000 xg for three times. The pellets were resuspended with 1.4x RE buffer without or with a hybridization oligo to restore the *EcoR*I restriction sites and fast frozen using dry ice, then stored at –80 °C.

The DNA were solubilized by incubating the cells with 0.1% SDS on 65 °C for 13 minutes. 1% Triton X-100 quenched the SDS. The DNA was digested by 20 U *EcoR*I at 37 °C for 1 hour. The restriction enzyme was deactivated by incubating the DNA with 1.5% SDS on 55 °C for 10 minutes. The cells were put back on ice, and the SDS was quenched by the addition of 6% Triton X-100. Ligation buffer (50 mM Tris-HCl pH 8.0, 10 mM MgCl_2_, 10 mM DTT, 2.5 μg/mL BSA, 1 mM ATP pH 8.0, 8 U T4 DNA ligase) was added to perform ligation reaction at 16 °C for 1 hour and 30 minutes. 25 μg/mL proteinase K was added to digest the enzymes at 65 °C for 30 minutes. DNA was extracted by adding phenol:chloroform:isoamyl alcohol and vortex. The upper water phase was moved and incubated with a tenth volume of sodium acetate and a volume of isopropanol at room temperature for 30 minutes and centrifuged at 21,130 xg, 4 °C for 10 minutes to get DNA precipitation. The DNA pellets were dried at 37 °C and dissolved by incubating with 1x TE buffer (10 mM Tris-Cl pH 8.0, 1 mM EDTA) at 37 °C for 1 hour. The DNA was used as qPCR template. DLC chimera content was calculated by [DLC amplification efficiency]^[-Cp_(DLC)_], and the intramolecular ligation product content was calculated by [intramolecular ligation amplification efficiency]^[-Cp_(ligation)_]. The final DLC signal was calculated by DLC chimera content/intramolecular ligation product content.

## Notes

### Competing Interest Statement

The authors have declared no competing interest.

## Works Cited

1. Kowalczykowski, S.C. (2015) An Overview of the Molecular Mechanisms of Recombinational DNA Repair. Cold Spring Harb Perspect Biol, 7.

2. Jasin, M. and Rothstein, R. (2013) Repair of strand breaks by homologous recombination. Cold Spring Harb Perspect Biol, 5, a012740.

3. San Filippo, J., Sung, P. and Klein, H. (2008) Mechanism of eukaryotic homologous recombination. Annu Rev Biochem, 77, 229–257.

4. Piazza, A., Bordelet, H., Dumont, A., Thierry, A., Savocco, J., Girard, F. and Koszul, R. (2021) Cohesin regulates homology search during recombinational DNA repair. Nat Cell Biol, 23, 1176–1186.

5. Dumont, A., Mendiboure, N., Savocco, J., Anani, L., Moreau, P., Thierry, A., Modolo, L., Jost, D. and Piazza, A. (2024) Mechanism of homology search expansion during recombinational DNA break repair in Saccharomyces cerevisiae. Mol Cell, 84, 3237–3253.e3236.

6. Haber, J.E. (2018) DNA Repair: The Search for Homology. Bioessays, 40, e1700229.

7. Renkawitz, J., Lademann, C.A., Kalocsay, M. and Jentsch, S. (2013) Monitoring homology search during DNA double-strand break repair in vivo. Mol Cell, 50, 261–272.

8. Bell, J.C. and Kowalczykowski, S.C. (2016) RecA: Regulation and Mechanism of a Molecular Search Engine. Trends Biochem Sci, 41, 491–507.

9. Wiktor, J., Gynnå, A.H., Leroy, P., Larsson, J., Coceano, G., Testa, I. and Elf, J. (2021) RecA finds homologous DNA by reduced dimensionality search. Nature, 597, 426–429.

10. Yang, H. and Pavletich, N.P. (2021) Insights into homology search from cryo-EM structures of RecA-DNA recombination intermediates. Curr Opin Genet Dev, 71, 188–194.

11. Danilowicz, C., Fu, J. and Prentiss, M. (2024) Insight into RecA-mediated repair of double strand breaks is provided by probing how contiguous heterology aaects recombination. J Biol Chem, 300, 107887.

12. Qi, Z., Redding, S., Lee, J.Y., Gibb, B., Kwon, Y., Niu, H., Gaines, W.A., Sung, P. and Greene, E.C. (2015) DNA sequence alignment by microhomology sampling during homologous recombination. Cell, 160, 856–869.

13. Lee, J.Y., Terakawa, T., Qi, Z., Steinfeld, J.B., Redding, S., Kwon, Y., Gaines, W.A., Zhao, W., Sung, P. and Greene, E.C. (2015) DNA RECOMBINATION. Base triplet stepping by the Rad51/RecA family of recombinases. Science, 349, 977–981.

14. Greene, E.C. (2016) DNA Sequence Alignment during Homologous Recombination. J Biol Chem, 291, 11572–11580.

15. Ragunathan, K., Liu, C. and Ha, T. (2012) RecA filament sliding on DNA facilitates homology search. Elife, 1, e00067.

16. Forget, A.L. and Kowalczykowski, S.C. (2012) Single-molecule imaging of DNA pairing by RecA reveals a three-dimensional homology search. Nature, 482, 423–427.

17. Yang, H., Zhou, C., Dhar, A. and Pavletich, N.P. (2020) Mechanism of strand exchange from RecA-DNA synaptic and D-loop structures. Nature, 586, 801–806.

18. Xu, J., Zhao, L., Xu, Y., Zhao, W., Sung, P. and Wang, H.W. (2017) Cryo-EM structures of human RAD51 recombinase filaments during catalysis of DNA-strand exchange. Nat Struct Mol Biol, 24, 40–46.

19. Cloud, V., Chan, Y.L., Grubb, J., Budke, B. and Bishop, D.K. (2012) Rad51 is an accessory factor for Dmc1-mediated joint molecule formation during meiosis. Science, 337, 1222–1225.

20. De Vlaminck, I., van Loenhout, M.T., Zweifel, L., den Blanken, J., Hooning, K., Hage, S., Kerssemakers, J. and Dekker, C. (2012) Mechanism of homology recognition in DNA recombination from dual-molecule experiments. Mol Cell, 46, 616–624.

21. Xu, W., Dunlap, D. and Finzi, L. (2021) Energetics of twisted DNA topologies. Biophysical Journal, 120, 3242–3252.

22. Strick, T., Allemand, J.-F., Croquette, V. and Bensimon, D. (2000) Twisting and stretching single DNA molecules. Progress in Biophysics and Molecular Biology, 74, 115–140.

23. Sheinin, M.Y., Forth, S., Marko, J.F. and Wang, M.D. (2011) Underwound DNA under Tension: Structure, Elasticity, and Sequence-Dependent Behaviors. Physical Review Letters, 107, 108102.

24. Vlassakis, J., Feinstein, E., Yang, D., Tilloy, A., Weiller, D., Kates-Harbeck, J., Coljee, V. and Prentiss, M. (2013) Tension on dsDNA bound to ssDNA-RecA filaments may play an important role in driving eaicient and accurate homology recognition and strand exchange. Physical Review E, 87, 032702.

25. Danilowicz, C., Peacock-Villada, A., Vlassakis, J., Facon, A., Feinstein, E., Kleckner, N. and Prentiss, M. (2014) The diaerential extension in dsDNA bound to Rad51 filaments may play important roles in homology recognition and strand exchange. Nucleic Acids Res, 42, 526–533.

26. van der Heijden, T., Modesti, M., Hage, S., Kanaar, R., Wyman, C. and Dekker, C. (2008) Homologous recombination in real time: DNA strand exchange by RecA. Mol Cell, 30, 530–538.

27. Tavares, E.M., Wright, W.D., Heyer, W.D., Le Cam, E. and Dupaigne, P. (2019) In vitro role of Rad54 in Rad51-ssDNA filament-dependent homology search and synaptic complexes formation. Nat Commun, 10, 4058.

28. Solinger, J.A., Lutz, G., Sugiyama, T., Kowalczykowski, S.C. and Heyer, W.D. (2001) Rad54 protein stimulates heteroduplex DNA formation in the synaptic phase of DNA strand exchange via specific interactions with the presynaptic Rad51 nucleoprotein filament. J Mol Biol, 307, 1207–1221.

29. Crickard, J.B., Moevus, C.J., Kwon, Y., Sung, P. and Greene, E.C. (2020) Rad54 Drives ATP Hydrolysis-Dependent DNA Sequence Alignment during Homologous Recombination. Cell, 181, 1380–1394.e1318.

30. Sridalla, K., Woodhouse, M.V., Hu, J., Scheer, J., Ferlez, B. and Crickard, J.B. (2024) The translocation activity of Rad54 reduces crossover outcomes during homologous recombination. Nucleic Acids Res, 52, 7031–7048.

31. Wolner, B., van Komen, S., Sung, P. and Peterson, C.L. (2003) Recruitment of the recombinational repair machinery to a DNA double-strand break in yeast. Mol Cell, 12, 221–232.

32. Renkawitz, J., Lademann, C.A. and Jentsch, S. (2014) Mechanisms and principles of homology search during recombination. Nat Rev Mol Cell Biol, 15, 369–383.

33. Flaus, A., Martin, D.M., Barton, G.J. and Owen-Hughes, T. (2006) Identification of multiple distinct Snf2 subfamilies with conserved structural motifs. Nucleic Acids Res, 34, 2887–2905.

34. Wolner, B. and Peterson, C.L. (2005) ATP-dependent and ATP-independent Roles for the Rad54 Chromatin Remodeling Enzyme during Recombinational Repair of a DNA Double Strand Break*. Journal of Biological Chemistry, 280, 10855–10860.

35. Zhang, Z., Fan, H.Y., Goldman, J.A. and Kingston, R.E. (2007) Homology-driven chromatin remodeling by human RAD54. Nat Struct Mol Biol, 14, 397–405.

36. Alexeev, A., Mazin, A. and Kowalczykowski, S.C. (2003) Rad54 protein possesses chromatin-remodeling activity stimulated by the Rad51-ssDNA nucleoprotein filament. Nat Struct Biol, 10, 182–186.

37. Amitani, I., Baskin, R.J. and Kowalczykowski, S.C. (2006) Visualization of Rad54, a chromatin remodeling protein, translocating on single DNA molecules. Mol Cell, 23, 143–148.

38. Thomä, N.H., Czyzewski, B.K., Alexeev, A.A., Mazin, A.V., Kowalczykowski, S.C. and Pavletich, N.P. (2005) Structure of the SWI2/SNF2 chromatin-remodeling domain of eukaryotic Rad54. Nat Struct Mol Biol, 12, 350–356.

39. Hopfner, K.P., Gerhold, C.B., Lakomek, K. and Wollmann, P. (2012) Swi2/Snf2 remodelers: hybrid views on hybrid molecular machines. Curr Opin Struct Biol, 22, 225–233.

40. Mazin, A.V., Bornarth, C.J., Solinger, J.A., Heyer, W.D. and Kowalczykowski, S.C. (2000) Rad54 protein is targeted to pairing loci by the Rad51 nucleoprotein filament. Mol Cell, 6, 583–592.

41. Petukhova, G., Van Komen, S., Vergano, S., Klein, H. and Sung, P. (1999) Yeast Rad54 promotes Rad51-dependent homologous DNA pairing via ATP hydrolysis-driven change in DNA double helix conformation. J Biol Chem, 274, 29453–29462.

42. Petukhova, G., Stratton, S. and Sung, P. (1998) Catalysis of homologous DNA pairing by yeast Rad51 and Rad54 proteins. Nature, 393, 91–94.

43. Raschle, M., Van Komen, S., Chi, P., Ellenberger, T. and Sung, P. (2004) Multiple interactions with the Rad51 recombinase govern the homologous recombination function of Rad54. J Biol Chem, 279, 51973–51980.

44. Alexiadis, V., Lusser, A. and Kadonaga, J.T. (2004) A conserved N-terminal motif in Rad54 is important for chromatin remodeling and homologous strand pairing. J Biol Chem, 279, 27824–27829.

45. Crickard, J.B., Kwon, Y., Sung, P. and Greene, E.C. (2020) Rad54 and Rdh54 occupy spatially and functionally distinct sites within the Rad51-ssDNA presynaptic complex. Embo j, 39, e105705.

46. Wright, W.D. and Heyer, W.D. (2014) Rad54 functions as a heteroduplex DNA pump modulated by its DNA substrates and Rad51 during D loop formation. Mol Cell, 53, 420–432.

47. Shah, P.P., Zheng, X., Epshtein, A., Carey, J.N., Bishop, D.K. and Klein, H.L. (2010) Swi2/Snf2-related translocases prevent accumulation of toxic Rad51 complexes during mitotic growth. Mol Cell, 39, 862–872.

48. Mason, J.M., Dusad, K., Wright, W.D., Grubb, J., Budke, B., Heyer, W.D., Connell, P.P., Weichselbaum, R.R. and Bishop, D.K. (2015) RAD54 family translocases counter genotoxic eaects of RAD51 in human tumor cells. Nucleic Acids Res, 43, 3180–3196.

49. Li, X., Zhang, X.P., Solinger, J.A., Kiianitsa, K., Yu, X., Egelman, E.H. and Heyer, W.D. (2007) Rad51 and Rad54 ATPase activities are both required to modulate Rad51-dsDNA filament dynamics. Nucleic Acids Res, 35, 4124–4140.

50. Kiianitsa, K., Solinger, J.A. and Heyer, W.D. (2006) Terminal association of Rad54 protein with the Rad51-dsDNA filament. Proc Natl Acad Sci U S A, 103, 9767–9772.

51. Van Komen, S., Petukhova, G., Sigurdsson, S., Stratton, S. and Sung, P. (2000) Superhelicity-driven homologous DNA pairing by yeast recombination factors Rad51 and Rad54. Mol Cell, 6, 563–572.

52. Crickard, J.B. and Greene, E.C. (2019) Helicase Mechanisms During Homologous Recombination in Saccharomyces cerevisiae. Annu Rev Biophys, 48, 255–273.

53. Ceballos, S.J. and Heyer, W.D. (2011) Functions of the Snf2/Swi2 family Rad54 motor protein in homologous recombination. Biochim Biophys Acta, 1809, 509–523.

54. Ristic, D., Wyman, C., Paulusma, C. and Kanaar, R. (2001) The architecture of the human Rad54–DNA complex provides evidence for protein translocation along DNA. Proceedings of the National Academy of Sciences, 98, 8454–8460.

55. Tan, T.L., Kanaar, R. and Wyman, C. (2003) Rad54, a Jack of all trades in homologous recombination. DNA Repair (Amst), 2, 787–794.

56. Sanchez, H., Kertokalio, A., van Rossum-Fikkert, S., Kanaar, R. and Wyman, C. (2013) Combined optical and topographic imaging reveals diaerent arrangements of human RAD54 with presynaptic and postsynaptic RAD51-DNA filaments. Proc Natl Acad Sci U S A, 110, 11385–11390.

57. Prasad, T.K., Robertson, R.B., Visnapuu, M.L., Chi, P., Sung, P. and Greene, E.C. (2007) A DNA-translocating Snf2 molecular motor: Saccharomyces cerevisiae Rdh54 displays processive translocation and extrudes DNA loops. J Mol Biol, 369, 940–953.

58. Blumberg, S., Tkachenko, A.V. and Meiners, J.-C. (2005) Disruption of Protein-Mediated DNA Looping by Tension in the Substrate DNA. Biophysical Journal, 88, 1692–1701.

59. Deveryshetty, J., Mistry, A., Pangeni, S., Ghoneim, M., Tokmina-Lukaszewska, M., Kaushik, V., Taddei, A., Ha, T., Bothner, B. and Antony, E. (2024) Rad52 sorts and stacks Rad51 at the DNA junction to promote homologous recombination. bioRxiv.

60. Baumann, C.G., Bloomfield, V.A., Smith, S.B., Bustamante, C., Wang, M.D. and Block, S.M. (2000) Stretching of Single Collapsed DNA Molecules. Biophysical Journal, 78, 1965–1978.

61. Le, T.T., Wu, M., Lee, J.H., Bhatt, N., Inman, J.T., Berger, J.M. and Wang, M.D. (2023) Etoposide promotes DNA loop trapping and barrier formation by topoisomerase II. Nat Chem Biol, 19, 641–650.

62. Forth, S., Deufel, C., Sheinin, M.Y., Daniels, B., Sethna, J.P. and Wang, M.D. (2008) Abrupt Buckling Transition Observed during the Plectoneme Formation of Individual DNA Molecules. Physical Review Letters, 100, 148301.

63. Lee, J., Wu, M., Inman, J.T., Singh, G., Park, S.h., Lee, J.H., Fulbright, R.M., Hong, Y., Jeong, J., Berger, J.M., et al. (2023) Chromatinization modulates topoisomerase II processivity. Nature Communications, 14, 6844.

64. Piazza, A., Shah, S.S., Wright, W.D., Gore, S.K., Koszul, R. and Heyer, W.D. (2019) Dynamic Processing of Displacement Loops during Recombinational DNA Repair. Mol Cell, 73, 1255–1266.e1254.

65. Reitz, D., Savocco, J., Piazza, A. and Heyer, W.D. (2022) Detection of Homologous Recombination Intermediates via Proximity Ligation and Quantitative PCR in Saccharomyces cerevisiae. J Vis Exp.

66. Ding, Y., Manzo, C., Fulcrand, G., Leng, F., Dunlap, D. and Finzi, L. (2014) DNA supercoiling: A regulatory signal for the λ repressor. Proceedings of the National Academy of Sciences, 111, 15402–15407.

67. Fogg, J.M., Judge, A.K., Stricker, E., Chan, H.L. and Zechiedrich, L. (2021) Supercoiling and looping promote DNA base accessibility and coordination among distant sites. Nature Communications, 12, 5683.

68. Bianco, P.R., Bradfield, J.J., Castanza, L.R. and Donnelly, A.N. (2007) Rad54 oligomers translocate and cross-bridge double-stranded DNA to stimulate synapsis. J Mol Biol, 374, 618–640.

69. Liu, W., Saito, Y., Jackson, J., Bhowmick, R., Kanemaki, M.T., Vindigni, A. and Cortez, D. (2023) RAD51 bypasses the CMG helicase to promote replication fork reversal. Science, 380, 382–387.

70. Liu, S., Miné-Hattab, J., Villemeur, M., Guerois, R., Pinholt, H.D., Mirny, L.A. and Taddei, A. (2023) In vivo tracking of functionally tagged Rad51 unveils a robust strategy of homology search. Nat Struct Mol Biol, 30, 1582–1591.

71. Friskes, A., Snoek, M., Oldenkamp, R., Broek, B.v.d., Nahidiazar, L., Koob, L., Dick, A.E., Mertz, M., Harkes, R., Rowland, B.D., et al. (2025) Visualizing homology search in living cells. bioRxiv, 2025.2003.2001.640932.

72. Belan, O., Greenhough, L., Kuhlen, L., Anand, R., Kaczmarczyk, A., Gruszka, D.T., Yardimci, H., Zhang, X., Rueda, D.S., West, S.C. et al. (2023) Visualization of direct and diausion-assisted RAD51 nucleation by full-length human BRCA2 protein. Mol Cell, 83, 2925–2940.e2928.

73. Neal, F.E., Li, W., Uhrig, M.E., Katz, J.N., Syed, S., Sharma, N., Dutta, A., Burma, S., Hromas, R., Mazin, A.V. et al. (2025) Distinct roles of the two BRCA2 DNA-binding domains in DNA damage repair and replication fork preservation. Cell Rep, 44, 115654.

74. Pires, E., Sung, P. and Wiese, C. (2017) Role of RAD51AP1 in homologous recombination DNA repair and carcinogenesis. DNA Repair (Amst), 59, 76–81.

75. Selemenakis, P., Sharma, N., Uhrig, M.E., Katz, J., Kwon, Y., Sung, P. and Wiese, C. (2022) RAD51AP1 and RAD54L Can Underpin Two Distinct RAD51-Dependent Routes of DNA Damage Repair via Homologous Recombination. Front Cell Dev Biol, 10, 866601.

76. Uhrig, M.E., Sharma, N., Maxwell, P., Gomez, J., Selemenakis, P., Mazin, A.V. and Wiese, C. (2024) Disparate requirements for RAD54L in replication fork reversal. Nucleic Acids Res, 52, 12390–12404.

77. Le, T.T., Gao, X., Park, S.H., Lee, J., Inman, J.T., Lee, J.H., Killian, J.L., Badman, R.P., Berger, J.M. and Wang, M.D. (2019) Synergistic Coordination of Chromatin Torsional Mechanics and Topoisomerase Activity. Cell, 179, 619–631.e615.

