## Supplementary material for "The Eukaryotic homology search complex distorts donor DNA structure to probe for homology": N/A

### Supplemental Information:

**Supplementary Table 1**

Strains used in the study

| Strains | Genotype | Source or reference |
|---|---|---|
| WDHY5511 | *ura3::A-HOcs, lys2::A, trp1::GAL-HO-hphMX, his3D200, can1-100, leu2-3,112, ade2-1, RAD5.* | Piazza et al, *Methods in Enzymology*, 2018 |
| WDHY5511 | *rad54Δ::KANMX* | This study |
| WDHY5511 | *RAD54::KANMX* | This study |
| WDHY5511 | *rad54 R272Q::KANMX* | This study |
| WDHY5511 | *rad54 R272A::KANMX* | This study |
| Protease deficient yeast strain | *MAT alpha leu2 trp1 ura3–52 prb1–1122 his3::pGAL1/10* | Crickard et al, *Cell*, 2020 |

**Supplementary Table 2**

Plasmid in the study

| Backbone | Construction | Source |
|---|---|---|
| pRS305 | *RAD54-KANMX* | This study |
| pRS305 | *rad54 R272A-KANMX* | This study |
| pRS305 | *rad54 R272Q-KANMX* | This study |
| pYES | *GST-GFP-RAD54* | Crickard et al EMBO 2018 |
| pYES | *GST-GFP-RAD54 R272Q* | Sridalla et al Nucleic Acid Research 2024 |
| pYES | *GST-GFP-RAD54 R272A* | Sridalla et al Nucleic Acid Research 2024 |
| pET11C | *6xHis-SUMO-yRad51* | Crickard et al Cell 2020 |
| pET11C | *6xHis-SUMO-yRad51IIA* | This Study |
| pNRFTC (pMDW111) | For torsionally constrained DNA | Le et al Cell 2019 |

**Supplementary Table 3**

Oligos used in the study

| Name | Sequence 5' to 3' | Purpose |
|---|---|---|
| 90-mer DNA | Atto647N-GATGTTCTGCTGGATATGCACTTTTCCGGGCTGACGTACACCGTGCTCAGCCTGTTTTTCAGCGATCCGGATATGCATCCGCTGGATTTC | Single Molecule imaging |
| pUC57-F | GTAAAACGACGGCCAGTG | To make Biotin and digoxigenin labeled adapters (Le et al Cell 2019) |
| pUC57-R | GGAAACAGCTATGACCATG | To make Biotin and digoxigenin labeled adapters (Le et al Cell 2019) |
| olWDH1760 | CAGCGGGCTTGCAGAAGTTG | To amplify genomic DNA at *ARG4* |
| olWDH1761 | GGCCAATTAGTTCACCAAGACG | To amplify genomic DNA at *ARG4* |
| olWDH1766 | GTTTCAGCTTTCCGCAACAG | To quantify DSB induction |

| | | |
|---|---|---|
| olWDH1767 | GGCGAGGTATTGGATAGTTCC | To quantify DSB induction |
| olWDH1762 | ACTTCGAATTTCGGCACTTC | To quantify intramolecular ligation efficiency of *Eco*RI-derived fragments |
| olWDH1763 | CGATGAAACGTTAAGTGACCAC | To quantify intramolecular ligation efficiency of *Eco*RI-derived fragments |
| olWDH1764 | AGAGCGGTCAGTAGCAATCC | To amplify at the upstream of DSB |
| olWDH1765 | CACACGCGAAAAACCGCC | To amplify at the upstream of the donor DNA, used with olWDH1764 to quantify DLC signal |
| olWDH2019 | CTTTAACCGGACGCTCGA | To quantify psoralen crosslinking efficiency |
| olWDH2020 | TTGAGTTTATTGCTGCCGTC | To quantify psoralen crosslinking efficiency |
| olWDH1768 | AGGAGCACAGACTTAGATTGG | Used with olWDH1764 to measure *Eco*RI recognition site restoration |
| olWDH1770 | CGAAATCATCTTCGGTTAAATCCAAAACGGCAGAAGCCTGAATGAAACATATGAACCAATTGGAGGACGTCAATGAATTCTGGGATCCATTGCATTTTT | To restore *Eco*RI site |

### Supplemental Figure 1

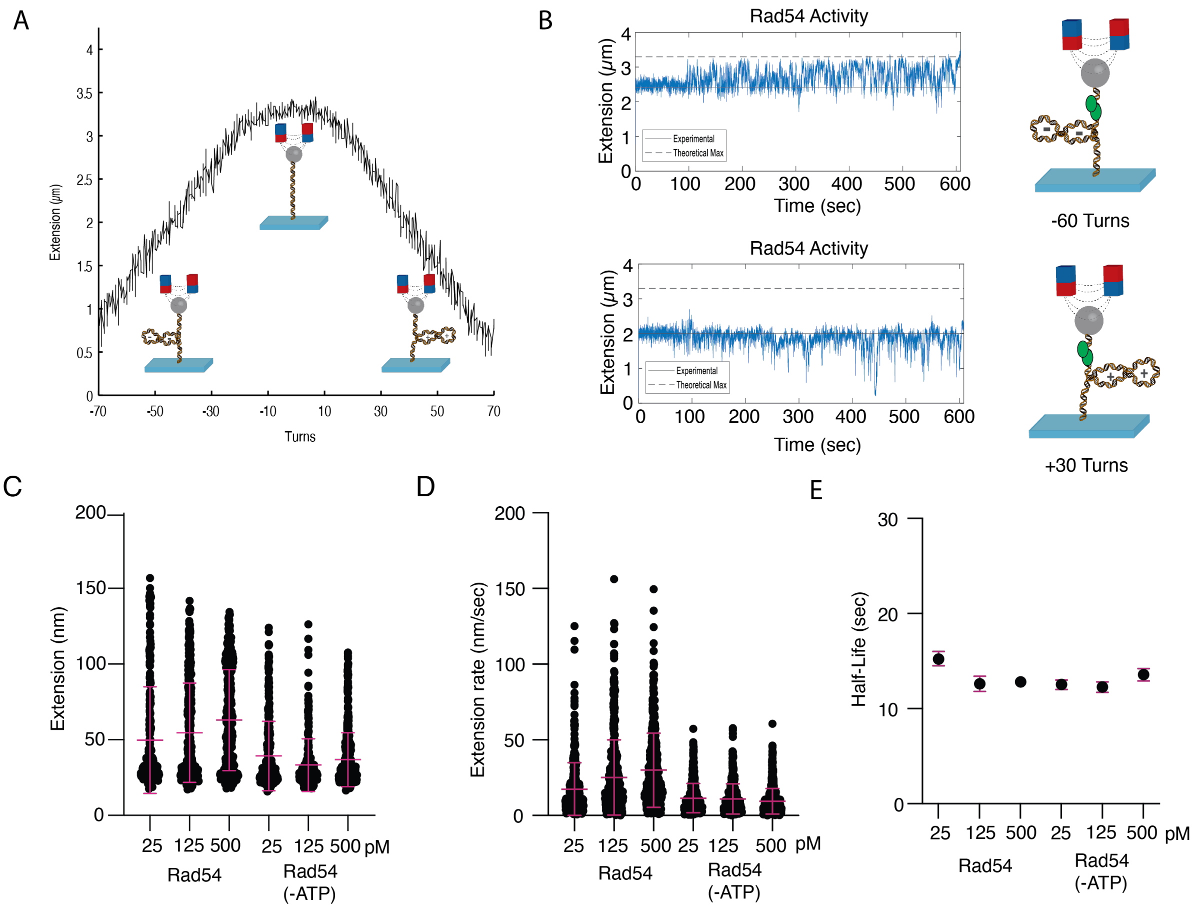


**Supplemental Figure 1: Rad54 enhances and stabilizes longer stretches of underwound DNA (A).** Example hat curve that illustrates the state of the DNA within each curve regime. At -60, the DNA forms a negative plectoneme, and at +30, a positive plectoneme. DNA at the apex is extended DNA. **(B).** Examples of Rad54 activity in the hat curve's -60 (Top) and +30 (Bottom) regions. The dashed line represents the max extension **(C).** Dot plot representing the extension of Rad54 +ATP at 25 (N=262), 125 (N=351), and 500 pM (N=350) and Rad54 -ATP at 25 (N=297), 125 (N=272), and 500 pM(N=466). The bar represents the mean, and the error bars represent the standard deviation. **(D).** Dot plot representing the extension per sec for Rad54 +ATP at 25 (N=388), 125 (N=462), and 500 pM (N=438) and Rad54 -ATP at 25 (N=464), 125 (N=409), 500 pM (N=693). The bar represents the mean, and the error bars represent the standard deviation of the data. **(E).** Graph representing the half-life of Rad54 at 25, 125, and 500 pM with and without ATP. The dots represent the half-life, and the bars represent the 95% confidence of the fit.

### Supplemental Figure 2

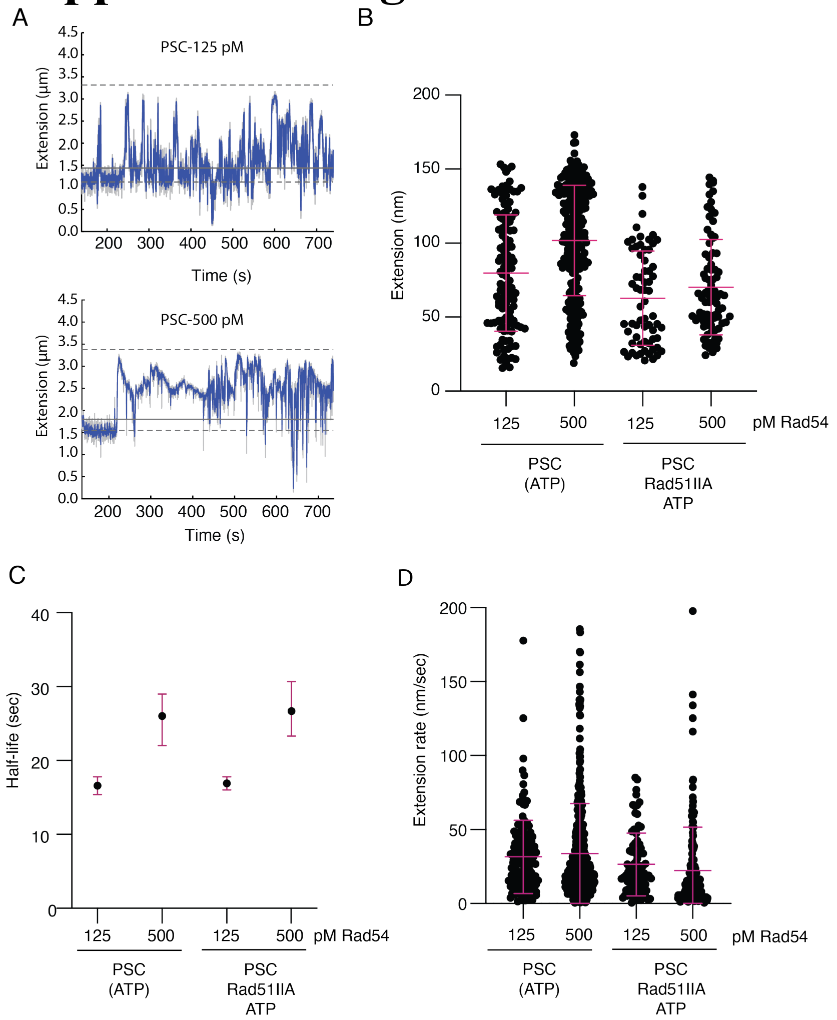


**Supplemental Figure 2: Impact of concentration on PSC activity**

**(A).** Representative traces for magnetic tweezer activity monitor for DNA molecules starting at -60 turns. The traces represent 125 pM (Top) and 500 pM (Bottom) PSC conditions. The top dashed line is the max extension, and the bottom dashed line is the DNA baseline. The solid line is the 3 standard deviation used as the cut off for analysis. **(B).** Dot plot representing the extension added for PSCs at 125 pM (N=117) and 500 pM (N=277), and PSC with Rad51IIA at 125 pM (N=65) and 500 pM (N=82). The bar represents the mean of the data, and the error bars represent the standard deviation. **(C).** Dot plot representing the extension per second for PSC 125 pM (N=148) and 500 pM (N=443), and PSC with Rad51IIA at 125 pM (N=84) and 500 pM (N=161). The bar represents the mean of the data, and the error bars represent the standard deviation of the data. **(D).** Graph representing the Half-life measurements for PSC at 125 pM (N=120) and 500 pM (N=277), and PSC with Rad51IIA at 125 pM (N=65) and 500 pM (N=97). The dot represents the half-life, and the error bars represent the 95% confidence interval for the fit.

### Supplemental Figure 3

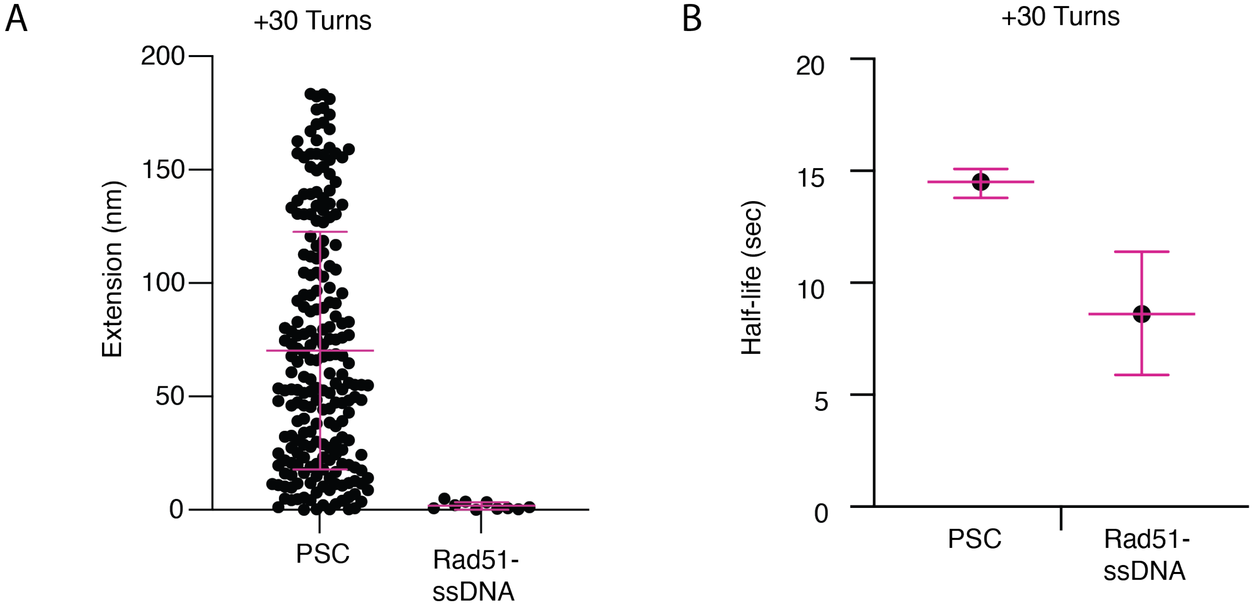


**Supplemental Figure 3: Rad54 significantly improves Rad51-ssDNA binding at +30 turns (A).** Extension measurements for PSC (N=215) and Rad51-ssDNA alone (N=10). The bar represents the mean, and the error bars represent the standard deviation of the data. **(B).** Half-life measurements for extension events for PSC (N=244) and Rad51-ssDNA alone (N=28). The dot represents the mean of the data, and the error bars represent the confidence in the fit.

### Supplemental Figure 4

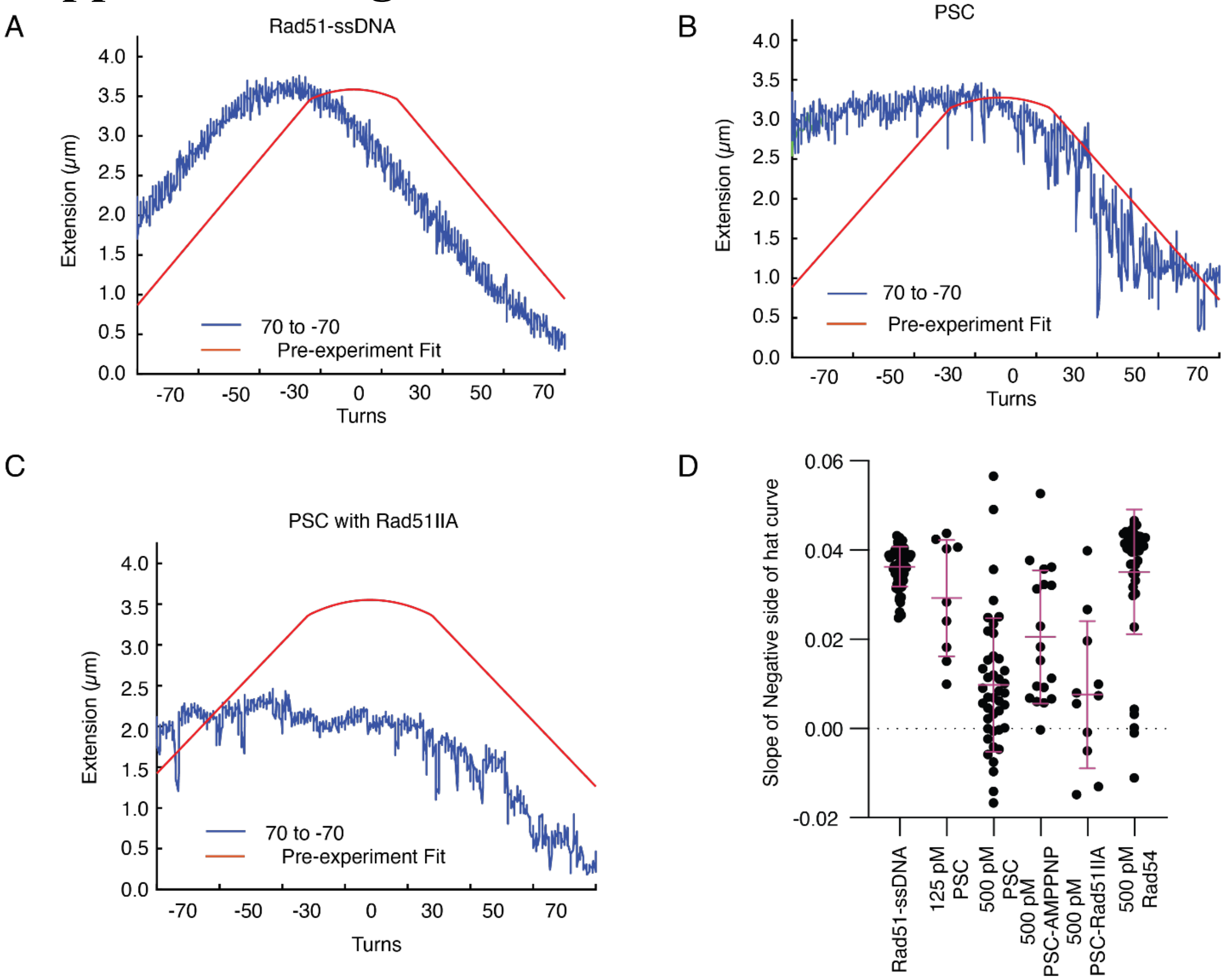


**Supplemental Figure 4: Post experiment hat curves show additional force added by Rad54** **(A)** Representative post-experiment hat curve for Rad51-ssDNA alone. The pre-hat curve is in Red, and the post experiment curve is in blue. **(B).** Representative post-experiment hat curve for the PSC. The pre-hat curve is in Red, and the post-experiment curve is in blue. **(C).** Representative post-experiment hat curve for PSC with Rad51-IIA. The pre-hat curve is in Red, and the post-experiment curve is in blue. **(D).** Graph representing the slope of the -70 to -20 turns of the post-experiment hat curve for Rad51-ssDNA (N=59), PSC 125 pM (N=9), PSC 500 pM (N=42), PSC with AMPPNP (N=18), PSC with Rad51IIA 500 pM (N=11), and Rad54 alone 500 pM (N=44) The bar represents the mean, and the error bars the standard deviation of the data.

### Supplemental Figure 5

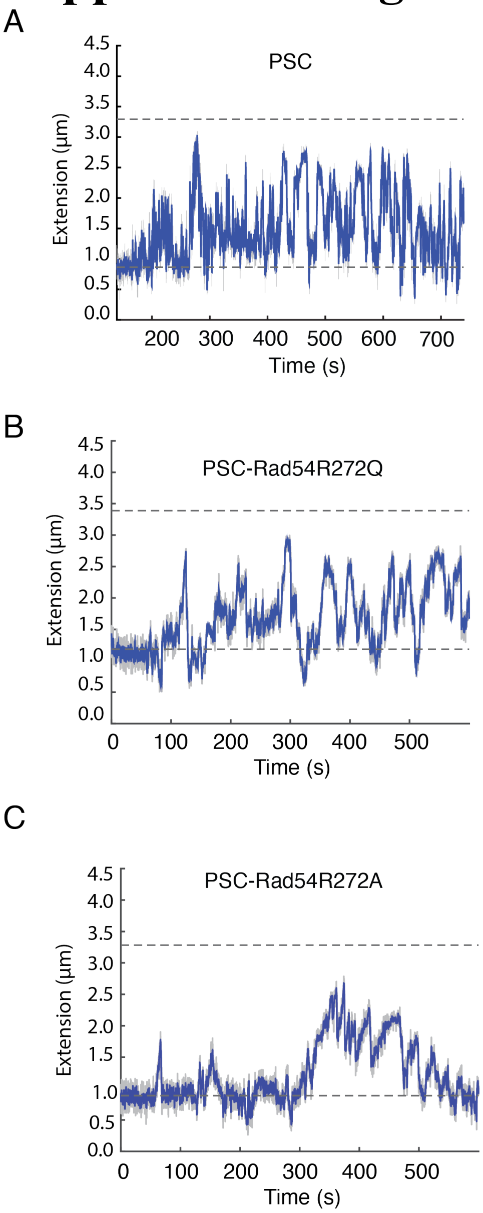


**Supplemental Figure 5: Representative traces for MT experiments at 125 pM**
**(A).** Representative MT trace for PSC at 125 pM. **(B).** Representative MT trace for PSC with Rad54 R272Q at 125 pM**. (C).** Representative MT trace for PSC with Rad54 R272A at 125 pM. The top dashed line represents max DNA extension, and the bottom dashed line represents the DNA baseline.

### Supplemental Figure 6

Rad51 and Rad54 global homology search

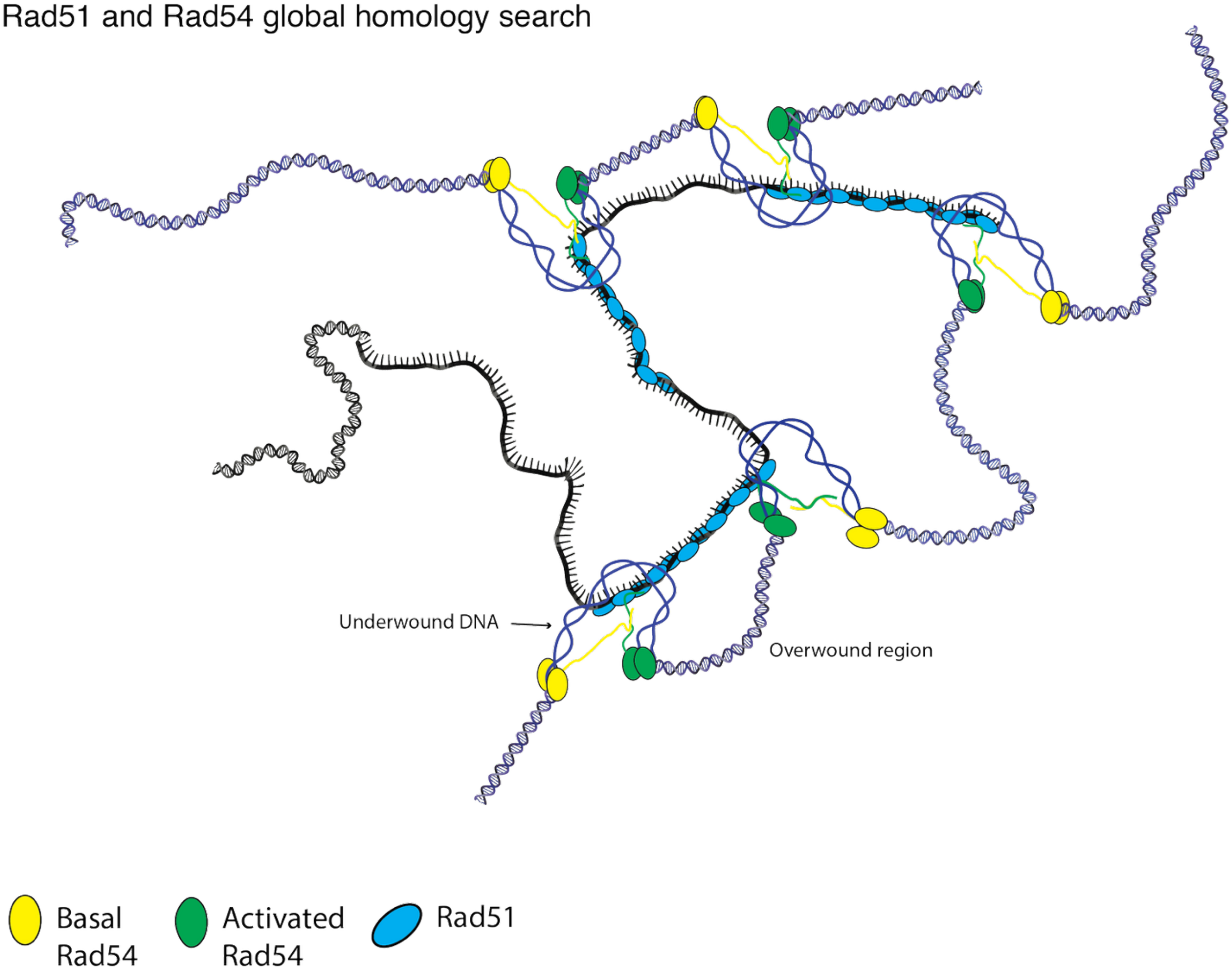


**Supplemental Figure 6: Model for multi-domain homology search**
A model representing a potential mechanism for multi domain homology search.
